# Switching Neuron Contributions to Second Network Activity

**DOI:** 10.1101/2023.10.05.560974

**Authors:** Savanna-Rae H Fahoum, Dawn M Blitz

**Affiliations:** Department of Biology and Center for Neuroscience, Miami University, Oxford OH 45056

**Keywords:** Neuronal switching, central pattern generator, neuromodulation, coordination, degeneracy

## Abstract

Network flexibility is important for adaptable behaviors. This includes neuronal switching, where neurons alter their network participation, including changing from single-to dual-network activity. Understanding the implications of neuronal switching requires determining how a switching neuron interacts with each of its networks. Here, we tested 1) whether “home” and second networks, operating via divergent rhythm generation mechanisms, regulate a switching neuron, and 2) if a switching neuron, recruited via modulation of intrinsic properties, contributes to rhythm or pattern generation in a new network. Small, well-characterized feeding-related networks (pyloric, ∼1 Hz; gastric mill, ∼0.1 Hz) and identified modulatory inputs make the isolated crab (*Cancer borealis*) stomatogastric nervous system (STNS) a useful model to study neuronal switching. In particular, the neuropeptide Gly^1^-SIFamide switches the lateral posterior gastric (LPG) neuron (2 copies) from pyloric-only to dual-frequency pyloric/gastric mill (fast/slow) activity via modulation of LPG intrinsic properties. Using current injections to manipulate neuronal activity, we found that gastric mill, but not pyloric, network neurons regulated the intrinsically generated LPG slow bursting. Conversely, selective elimination of LPG from both networks using photoinactivation revealed that LPG regulated gastric mill neuron firing frequencies but was not necessary for gastric mill rhythm generation or coordination. However, LPG alone was sufficient to produce a distinct pattern of network coordination. Thus, modulated intrinsic properties underlying dual-network participation may constrain which networks can regulate switching neuron activity. Further, recruitment via intrinsic properties may occur in modulatory states where it is important for the switching neuron to actively contribute to network output.

**New and Noteworthy:** We used small, well-characterized networks to investigate interactions between rhythmic networks and neurons that switch their network participation. For a neuron switching into dual-network activity, only the second network regulated its activity in that network. Additionally, the switching neuron was sufficient but not necessary to coordinate second network neurons, and regulated their activity levels. Thus, regulation of switching neurons may be selective, and a switching neuron is not necessarily simply a follower in additional networks.

## Introduction

Oscillatory networks, including central pattern generator (CPG) networks generating rhythmic patterns such as walking, breathing, and chewing are highly modulated to promote adaptation (1–9). CPG flexibility includes neuronal switching, where neuronal participation changes from one network to another, or to multiple networks, via neuromodulation of intrinsic and/or synaptic properties (10–17). Although CPG neurons are typically involved in rhythm and/or pattern generation in their home network, it is unclear whether switching neurons from other networks also take on such an active role. For instance, they may be passive followers of a new pattern imposed upon them or become active contributors to rhythm and/or pattern generation in the new network.

The same or different neuron complements can be involved in rhythm generation, producing rhythmic activity and setting rhythm frequency, and pattern generation, determining the relative timing among network neurons (6, 18–21). In most examples with identified cellular-level mechanisms, switching neurons are recruited into another network via modulation of inter-network synapses (10, 12, 14), and passively follow this input, without contributing to network output, such as rhythm frequency or network neuron coordination. However, neurons may also be recruited into a second network via modulation of intrinsic properties (15, 17, 22, 23). Here we ask whether a switching neuron has an active role in a novel network when the recruitment mechanism is intrinsic to the switching neuron.

Due to their smaller numbers of identified, accessible rhythm and pattern generator neurons with characterized connectomes, plus identified modulatory inputs, invertebrate systems are particularly useful for studying neuronal switching (24–32). In the crustacean stomatogastric nervous system (STNS), 26-30 neurons generate the pyloric (food filtering, ∼ 1 Hz) and gastric mill (food chewing, ∼ 0.1 Hz) rhythms (28, 29, 33–35). Furthermore, neuronal switching mechanisms are established in the STNS (10, 12, 17). Here, we examined whether a neuron recruited into dual-network activity via modulation of intrinsic properties contributes to rhythm and/or pattern generation in a second network.

In the crab, *Cancer borealis*, activation of the modulatory projection neuron 5 (MCN5), or bath application of the MCN5 neuropeptide Gly^1^-SIFamide, increases pyloric frequency (26), activates the gastric mill rhythm (36), and switches the pyloric-only lateral posterior gastric (LPG) neuron (2 copies) into dual-frequency pyloric/gastric mill bursting (17). During the MCN5/Gly^1^-SIFamide-elicited gastric mill rhythm, LPG is coordinated with the lateral gastric (LG), inferior cardiac (IC), and dorsal gastric (DG) network neurons (17, 36). While LPG slow bursting occurs via Gly^1^-SIFamide modulation of LPG intrinsic properties (17, 23), how LPG is incorporated into the gastric mill network, and whether it contributes to rhythm and/or pattern generation is unknown. We hypothesized that LPG actively contributes to rhythm and pattern generation of the Gly^1^-SIFamide gastric mill rhythm. Our results suggest that even if a switching neuron is not necessary for rhythm generation, it can contribute to pattern generation, shaping the activity strength and timing of other neurons in a second network.

## Materials and Methods

### Animals

Male *Cancer borealis* crabs were obtained from The Fresh Lobster company (Gloucester, MA) and maintained in tanks containing artificial seawater at 10°C -12°C. For experiments, crabs were anesthetized on ice for 30-50 min before the foregut was removed from the animal during gross dissection, bisected, and pinned flat in a Sylgard 170-lined dish (Thermo Fisher Scientific). The STNS was dissected from the foregut during fine dissection and pinned in a Sylgard 184-lined petri dish (Thermo Fisher Scientific) (37). Throughout the dissection, the preparation was kept chilled in *C. borealis* physiological saline at 4°C.

### Solutions

*C. borealis* physiological saline was composed of the following (in mM): 440 NaCl, 26 MgCl2,13 CaCl2,11 KCl,10 Trizma base, 5 Maleic acid, pH 7.4-7.6. Squid internal electrode solution contained the following (in mM): 10 MgCl2, 400 Potassium D-gluconic acid, 10 HEPES, 15 NaSO4, 20 NaCl, pH 7.45 (38). Gly^1^-SIFamide (GYRKPPFNG-SIFamide, custom peptide synthesis: Genscript) (17, 36, 39–41) was dissolved in optima water (Thermo Fisher Scientific) at 10^-2^ M and aliquots were stored at -20°C until needed. Gly^1^-SIFamide aliquots were diluted in physiological saline at a final concentration of 5 µM. Picrotoxin (PTX) powder (Sigma Aldrich) was added directly to physiological saline at a final concentration of 10 µM and vigorously stirred for 45 min before use (PTX). Gly^1^-SIFamide aliquots were added directly to PTX in physiological saline (10 µM) at a final concentration of 5 µM (SIF:PTX).

### Electrophysiology

All preparations were continuously superfused with chilled *C. borealis* saline (8-10°C), or chilled saline containing Gly^1^-SIFamide and/or PTX, as indicated. For uninterrupted superfusion of the preparation, all solution changes were performed using a switching manifold. Extracellular nerve activity was recorded using a model 1700 A-M Systems Amplifier and custom-made stainless-steel pin electrodes. Vaseline wells were built around nerves to isolate neuron signals, with one stainless-steel electrode wire placed inside the well, and the other outside the well as reference. Stomatogastric ganglion (STG) somata were exposed by removing the thin layer of tissue across the ganglion and observed with light transmitted through a dark-field condenser (MBL-12010 Nikon Instruments). Intracellular recordings were collected using sharp-tip glass microelectrodes (18-40 MΩ) tip-filled with squid internal electrode solution (*see Solutions*). STG neurons were identified based on their nerve projection patterns and their synaptic interactions with other STG neurons. MCN5 activation experiments included MCN5 stimulation via intracellular current injection, or extracellular inferior oesophageal nerve (*ion*) stimulation (30 Hz, tonic), in which a portion of the MCN1 axon was photoinactivaed (see below) as it entered the STG to prevent MCN1 neurotransmission during *ion* stimulation (17). MCN5 activation experiments were reanalyzed from 17. All intracellular recordings were collected using AxoClamp 900A amplifiers in current-clamp mode (Molecular Devices). All experiments were conducted in the isolated STNS following transection of the inferior and superior oesophageal nerves (*ion* and *son*, respectively, Fig. 1A). Electrophysiology recordings were collected using data acquisition hardware (Micro1401; Cambridge Electronic Design), software (Spike2; ∼5 kHz sampling rate, Cambridge Electronic Design), and laboratory computer (Dell).

**Figure 1.**
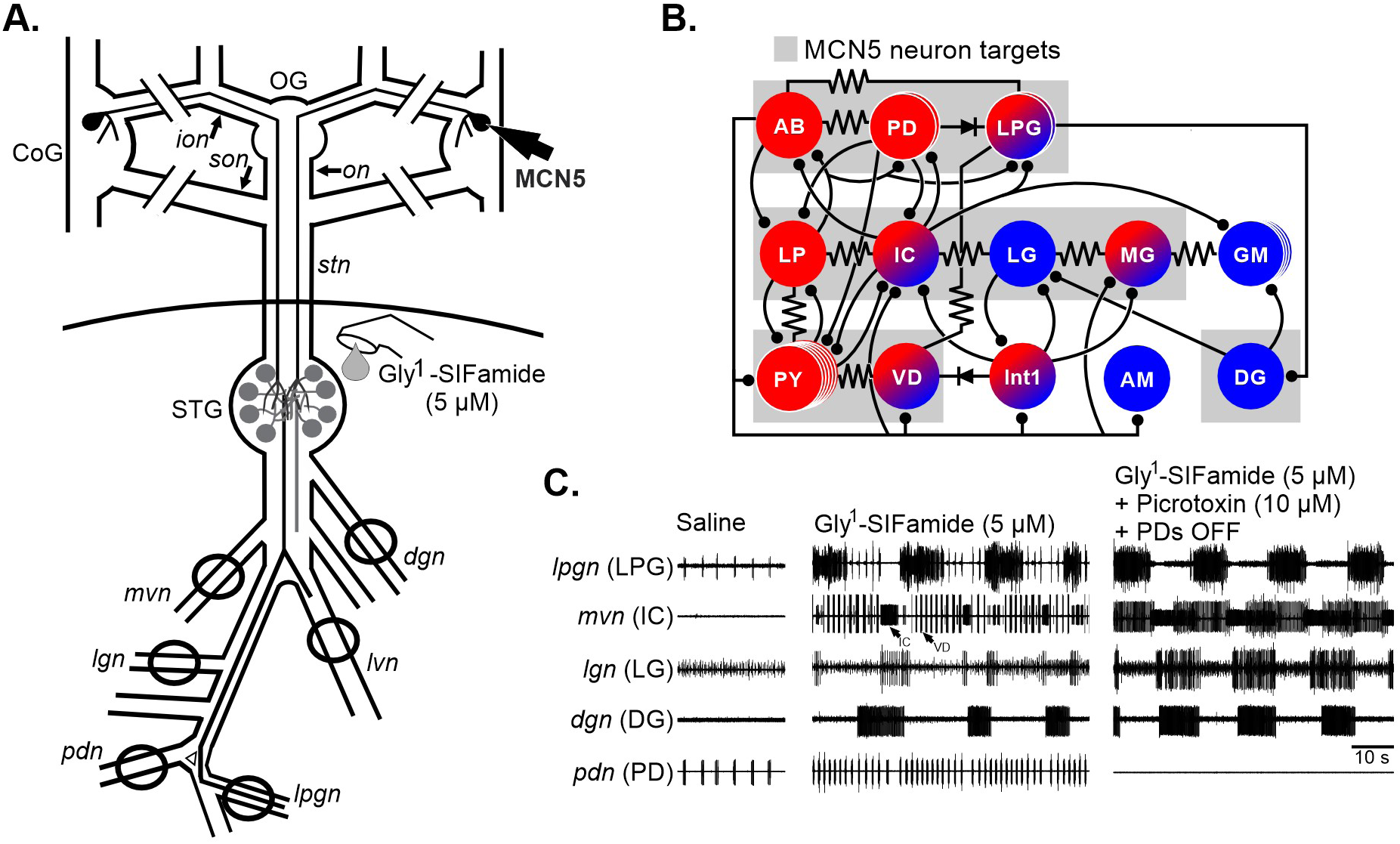
The isolated stomatogastric nervous system (STNS), including modulatory projection neuron MCN5, connectivity among the stomatogastric (STG) neurons, and STG gastric mill and pyloric response to bath application of the MCN5 neuropeptide Gly^1^-SIFamide. *A.* The isolated STNS is comprised of the paired commissural ganglia (CoG), oesophageal ganglion (OG), stomatogastric ganglion (STG), and connecting and peripheral nerves. The MCN5 neuron projects from the CoG through the *ion, on,* and *stn* to the STG (26). Hashed lines represent where *ions* and *sons* were cut to isolate STG neurons (see Materials and Methods). Black circles around nerves indicate Vaseline wells for extracellular recordings. Solid line across *stn* indicates a Vaseline wall which was made at the *stn* that extended across the dish to separate the STG and motor nerves from the modulatory CoGs, OG, and *ions* and *sons* during Gly^1^-SIFamide superfusion (liquid drop). ***B.*** Circuit diagram of the STG indicates chemical inhibitory (ball and stick) and electrical (resistor and diode symbols) synapses between pyloric (red), gastric mill (blue), and gastro-pyloric (red/blue circles) neurons. The MCN5 neuron acts on most STG neurons (grey backgrounds), including inhibiting pyloric neurons LP and PY, exciting the pyloric pacemaker ensemble (AB/PD/LPG), activating gastric mill neurons LG, IC, and DG, and switching pyloric-only LPG into dual pyloric/gastric mill timed bursting (17, 26, 36). ***C.*** *Left*, In baseline conditions with the *ions* and *sons* cut, only the pyloric rhythm is active (*Left*, Saline *lpgn* [LPG] and *pdn* [PD]). *Middle*, Bath application of MCN5 neuropeptide Gly^1^-SIFamide increases pyloric frequency (increased PD burst frequency), activates the gastric mill rhythm (*mvn* [IC], *lgn* [LG], *dgn* [DG]), and switches LPG into dual pyloric plus gastric mill-timed bursting. *Right*, In Gly^1^-SIFamide, when the LPG neuron is isolated from pyloric-timed input (PDs OFF) and gastric mill-timed input (PTX to block glutamatergic inhibition), LPG generates gastric mill-timed bursting intrinsically (17,23). Ganglia: CoG, commissural ganglion; OG, oesophageal ganglion; STG, stomatogastric ganglion. Nerves: *dgn*, dorsal gastric nerve; *ion*, inferior oesophageal nerve; *lgn*, lateral gastric nerve; *lpgn*, lateral posterior gastric nerve; *lvn*, lateral ventricular nerve; *mvn*, medial ventricular nerve; *on*, oesophageal nerve; *pdn*, pyloric dilator nerve; *son*, superior oesophageal nerve; *stn*, stomatogastric nerve; Neurons: AB, anterior burster; AM, anterior median; DG, dorsal gastric; GM, gastric mill; IC, inferior cardiac; Int1, interneuron 1; LG, lateral gastric; LP, lateral pyloric; LPG, lateral posterior gastric; MCN5, modulatory commissural neuron 5; MG, medial gastric; PD, pyloric dilator; PY, pyloric; VD, ventricular dilator.

Prior to all conducted experiments, unless otherwise indicated, the LP neuron was photoinactivated as described in 17. Briefly, the LP neuron was impaled with a sharp microelectrode (30-40 MΩ) that was tip-filled with AlexaFluor-568 hydrazide (10 mM in 200 mM KCl, Thermo Fisher Scientific) and back-filled with squid internal solution (*see Solutions*). The LP soma was filled using constant hyperpolarizing current injection (-5 nA) for 5-10 mins, and the current injection stopped to allow the dye to diffuse to the neurites and axon in the dorsal ventricular nerve (*dvn*) for 20-40 mins. The STG was then illuminated for 5-7 mins using a Texas red filter set (560 ± 40 nm wavelength; Leica Microsystems). Complete photoinactivation was confirmed when the LP membrane potential reached 0 mV. This same protocol was used for simultaneous photoinactivation of both copies of the LPG neuron (LPG Killed).

### LPG regulation via pyloric and gastric mill networks

To determine whether LPG gastric mill-timed (slow) bursting was regulated by the pyloric network in Gly^1^-SIFamide, we compared LPG bursting activity before, during, and after 200 s duration hyperpolarizing current injections into both pyloric dilator (PD) neurons. Due to electrical coupling between the two PD neurons and the anterior burster (AB), the pacemaker neuron for the pyloric rhythm, hyperpolarization of the PD neurons sufficiently hyperpolarizes the AB neuron as well, and thus eliminates the pyloric rhythm (17, 42). To regulate the pyloric rhythm frequency, steady depolarizing (0 to +2.5 nA) or hyperpolarizing (-6 to -0.25 nA) current was injected into the two PD neurons. Each current level was maintained for the duration of five LPG slow burst cycles before changing pyloric frequency with a new holding current. In an additional set of experiments, pyloric frequency was regulated via steady current injection while LPG was isolated from gastric mill synaptic input via PTX application (17). Briefly, PTX (10 µM) in physiological saline was bath applied to the preparation for a minimum of 15 min to allow for PTX to sufficiently block inhibitory glutamatergic synapses (43, 44). PTX was judged effective when activity of the IC and ventricular dilator (VD) neurons in the medial ventricular nerve (*mvn*) overlapped with one another or, in cases where the pyloric neuron lateral pyloric (LP) neuron was intact, glutamatergic inhibitory post-synaptic potentials from LP in the PD neuron were eliminated, and LP and pyloric (PY) neuron activity overlapped in the lateral ventricular nerve (*lvn*) (17, 43, 44). Gy^1^-SIFamide containing PTX (SIF:PTX, 5 µM:10 µM, respectively) was then applied to the preparation to examine the effects of pyloric frequency on LPG slow bursting with LPG isolated from glutamatergic synaptic inhibition.

To determine whether LPG slow burst activity was regulated by LG, IC, and/or DG gastric mill network neurons in Gly^1^-SIFamide application, we reanalyzed a dataset from our previous study (17), in which LG, IC, and DG neurons were hyperpolarized (-2 to -4 nA, 200 s) individually and simultaneously for 200 s, followed by 200 s with no manipulation. LPG slow burst duration (time from the first to the last action potential within a burst) and cycle period (time between the first action potential of two consecutive bursts) were measured 200 s before, during, and after each LG, IC, and DG hyperpolarization. To test whether LPG slow bursting regulated LG, IC, and DG gastric mill neuron activity, we measured LG, IC, and DG burst duration, cycle period, firing frequency ([number of spikes per burst – 1]/burst duration), and number of spikes per burst (the total count of action potentials between the first and last action potential of a burst) in Gly^1^-SIFamide conditions with LPG intact versus LPG Killed.

### Burst detection and identification

LPG neuron (2 copies) bursting activity was recorded in one or both of the lateral posterior gastric nerves (*lpgn*). LPG slow bursting was determined as described in 17. Briefly, a histogram containing LPG interspike intervals (ISIs) across an entire Gly^1^-SIFamide application was generated using a custom-written MATLAB script (MathWorks) and the two largest peaks identified. The first peak (within ∼0-0.5 s) included intra-burst intervals (interval between spikes during a burst), and the second peak (∼0.5-2 s) included inter-burst intervals (interval between spikes between bursts). The mean ISI between these two peaks was calculated and used as a cutoff value to identify LPG bursting, such that an ISI above the cutoff indicated the end of one burst and the beginning of another. To select only LPG slow bursts from all LPG bursts identified with the ISI cutoff, we used a custom written Spike2 script that identifies LPG bursts with a duration greater than one pyloric cycle period (from PD burst onset to the next PD burst onset, ∼1 s) (17).

To determine whether there was a relationship between pyloric frequency and LPG slow burst frequency, we measured LPG slow burst frequency (1/LPG slow burst cycle period) and burst duration in Gly^1^-SIFamide-only and SIF:PTX conditions (5 LPG slow burst cycles, 30-200 s time window). To examine the role of LPG slow bursting in LG, IC, and DG neuron regulation, we measured and compared LG, IC, and DG burst duration, cycle period, number of spikes per burst, and firing frequency across a 1200 s time window in Gly^1^-SIFamide with LPG Intact and LPG Killed conditions.

LG, IC, and DG bursting activity was recorded either intracellularly or via extracellular nerve recordings (*lgn, mvn, dgn*, respectively). All bursts had at least 3 action potentials with a maximum of 2 s interspike interval. IC pyloric-timed bursts were determined by including only IC bursts with a minimum duration of 0.45 s (36), each separated by the duration of a PD (*pdn*) burst. IC gastric mill-timed bursts were determined by grouping together consecutive pyloric-timed IC bursts with a minimum burst duration of 0.45 s, including up to one IC burst < 0.45 s in duration if it occurred between other IC bursts that were at least 0.45 s.

### Categorization of the gastric mill rhythm

To examine the role of LPG slow bursting in gastric mill network activity, we eliminated LPG actions by photoinactivating LPG (LPG Killed) with and without the influence of glutamatergic inhibitory synapses from network neurons LG, IC, and DG (PTX, 10 µM). To test whether LPG was necessary for generating the gastric mill rhythm, we used a combination of manipulations with 2-3 hr intervals between manipulations: Gly^1^-SIFamide with LPG Intact (SIF:LPG Intact); Gly^1^-SIFamide with LPG Killed (SIF:LPG Killed); Gly^1^-SIFamide with Picrotoxin (SIF:PTX); and SIF:PTX with LPG Killed (SIF:PTX LPG Killed). Not all manipulations were performed in all preparations. Specific comparisons included SIF:LPG Intact (LPG Included in the categorization analysis) versus SIF:LPG Intact (LPG Excluded from the categorization analysis) (n = 9, 41-142 cycles); SIF:LPG Intact (LPG Excluded) (n = 10, 41-142 cycles) versus SIF:LPG Killed (n = 10, 31-110 cycles); SIF:LPG Intact (LPG Included, n = 5, 29-105 cycles) versus SIF:PTX (LPG Included, n = 5, 52-114 cycles); and SIF:LPG Intact (LPG Excluded, n = 11, 29-105 cycles) versus SIF:PTX (LPG Excluded, n = 5, 52-114 cycles) versus SIF:PTX LPG Killed (n = 8, 8-124 cycles). For conditions without PTX application, the LP neuron was hyperpolarized to ensure LPG burst in gastric mill time (17). PTX was superfused prior to SIF:PTX, and PTX wash-out was confirmed by the presence of LP-elicited glutamatergic inhibitory post-synaptic potentials in an intracellular PD neuron recording (17, 43, 44).

The relative timing of neurons participating in the Gly^1^-SIFamide rhythm is variable. Therefore, we applied a qualitative cycle-by-cycle analysis using LG as a reference to characterize gastric mill network coordination (Fig. 5, method adapted from 45). First, LG, IC, DG, and LPG burst onsets and offsets were identified across a 1200 s time window in Spike2 and exported to MATLAB. IC, DG, and LPG burst onset and offset were compared relative to each LG cycle to determine the type of coordination for each neuron in each LG cycle. Then, the combination of coordination types was used to determine the network coordination among a total of four neurons (LG, IC, DG, and LPG) or three neurons (LG, IC, and DG), as indicated. For instance, complete coordination was defined as “All Coordinated” when all neurons assessed were associated with and defined as coordinated, with one cycle (Fig. 5). A neuron was considered to be “associated” with a current cycle if: burst onset and offset occurred within the current LG cycle; burst onset occurred near the end of the previous LG cycle (less than 10% into the previous cycle) plus burst offset occurring in the current LG cycle; or burst onset occurred in the current LG cycle (at least 10%) and burst offset in the next LG cycle. For instance, in the example traces shown for “All Coordinated” the DG neuron was identified as coordinated because its burst onset was greater than 10% in that LG cycle (Fig. 5Bi). If a neuron was active across a complete LG cycle and extended beyond that LG cycle, the cycle was classified as “One Neuron Tonic” (Fig. 5). We did not have instances of more than one neuron being tonic. When one or more neurons were silent, that cycle was defined as “One Neuron Silent” or “All Silent”, depending on the number not active in that cycle (Fig. 5). Finally, when IC, DG, and/or LPG neurons had more than one burst per LG cycle, or some other burst pattern that did not meet the above criteria, that cycle was defined as “One Neuron Uncoordinated” or “All Uncoordinated” (Fig. 5). Identified patterns from each preparation were accumulated and the mean percentage of cycles per category was calculated. To confirm that the results obtained were not due to multiple Gly^1^-SIFamide applications or due to photoinactivation itself, we performed control experiments where (1) LPG was intact for two consecutive Gly^1^-SIFamide applications and (2) two GM neurons were photoinactivated between two consecutive Gly^1^-SIFamide applications. All analyses were conducted after Gly^1^-SIFamide reached a steady state, approximately 10 mins after peptide wash-in.

### Spike phase analysis

To quantify the relative timing relationships of gastric mill neurons, all LG neuron cycles were used as reference, regardless of coordination type. Each LG cycle was divided into 100 bins and the number of spikes per bin quantified for all neurons, for all cycles, across the same 1200 s window used for categorical analysis. Action potentials and bursts were identified in Spike2 and exported to MATLAB. The number of spikes for each neuron, in each bin was accumulated using custom-written functions in MATLAB. Counts per bin were normalized to the total number of spikes in that neuron in each experiment, and expressed as a percent of total spikes.

### Software and statistical analysis

Raw electrophysiology data were analyzed using functions and scripts that were written in Spike2 or MATLAB. Statistical analyses were performed using SigmaPlot (Systat). Graphs were plotted in SigmaPlot or MATLAB and imported into CorelDraw. All final figures were created using CorelDraw (Corel). Data were analyzed for normality using the Shapiro-Wilk Normality test before determining whether to use a parametric or nonparametric test on each dataset. Pearson correlation, One-way repeated measures (RM) ANOVA, Friedmann’s one-way RM ANOVA on ranks, Paired t-test, Wilcoxon signed rank test, and *post hoc* tests for multiple comparisons were used as indicated. Overall p-values are reported in text, while *post-hoc* p-values are reported in tables, as indicated. Threshold for significance was p < 0.05. All data are presented as mean ± SEM.

### Code accessibility

Scripts and functions used for analysis were written in Spike2 and/or MATLAB and are available at https://github.com/blitzdm/Fahoum2023 or upon request from D.M.B.

## Results

The crab STNS is a region of the nervous system that controls the pyloric (food filtering, ∼1 Hz) and gastric mill (food chewing, ∼0.1 Hz) behaviors (Fig. 1) (28, 29). The STNS includes the paired commissural ganglia (CoG) and oesophageal ganglion (OG) containing modulatory projection neurons that have axonal projections to the STG (Fig. 1A). The STG is comprised of ∼30 neurons that form CPG networks driving the pyloric and gastric mill rhythms (Fig. 1B). These networks and their modulatory inputs are well-characterized, including identified synapses between neurons (Fig. 1B) (4, 29, 46). The neuropeptide Gly^1^-SIFamide is released from the modulatory projection neuron 5 (MCN5), which projects through the *ion* and stomatogastric nerve (*stn*) to the STG (Fig. 1A). Bath application or neuronal release of Gly^1^-SIFamide elicits an increase in pyloric frequency, activation of a gastric mill rhythm and switching of the pyloric-only LPG neuron (2 copies) into simultaneous dual pyloric/gastric mill timing (fast/slow timing, respectively) (Fig. 1C) (17, 26, 36). In saline conditions, LPG is pyloric-timed due to its electrical coupling with the pyloric pacemaker group (AB/PD/LPG) (Fig. 1C, left) (44, 47, S-RH Fahoum, DM Blitz, L Zhang, MP Nusbaum, unpublished). During Gly^1^-SIFamide application, LPG maintains its fast pyloric-timed bursting activity via electrical coupling, and additionally generates slow bursting in time with the gastric mill rhythm (Fig. 1C, middle). When LPG is isolated from pyloric and gastric mill inputs via hyperpolarization of the two PD neurons and application of picrotoxin (PTX, 10 µM), respectively, LPG generates slow, gastric mill-timed bursting independently due to Gly^1^-SIFamide modulation of LPG intrinsic properties (Fig. 1C, right) (17, 23). This mechanism of recruitment differs from other examples, in which modulation of synapses recruits a switching neuron into another network (10, 12, 13, 48).

During the MCN5/Gly^1^-SIFamide gastric mill rhythm, the LPG neuron is coordinated with the LG, IC, and DG network neurons (17, 36). Although Gly^1^-SIFamide modulation of LPG intrinsic properties is well-established as the mechanism for LPG to generate bursting at gastric mill rhythm frequency (17, 23), how LPG is incorporated into the gastric mill network, and whether it contributes to rhythm/pattern generation is not known. Here, we tested the hypothesis that LPG contributes to generating and shaping the Gly^1^-SIFamide gastric mill rhythm. First, we examined whether pyloric and gastric mill activity regulated the intrinsically-generated LPG gastric mill-timed (slow) bursting in Gly^1^-SIFamide. Then, we tested the role of LPG slow bursting in rhythm and/or pattern generation for the gastric mill network.

### Pyloric network regulation of LPG

In Gly^1^-SIFamide application, LPG periodically escapes its electrical coupling to the pyloric pacemaker group to burst in gastric mill time (17, 36). While this rhythmic electrical coupling input is not necessary for LPG to generate gastric mill-timed (slow) bursting, it is unknown whether the pyloric network regulates LPG slow bursting. In saline conditions, the AB neuron is the pacemaker for the pyloric rhythm, and its bursting activity controls rhythmic bursting in PD and LPG neurons through electrical coupling (47, 49, S-RH Fahoum, DM Blitz, L Zhang, MP Nusbaum, unpublished). We tested whether the pyloric network regulates LPG slow bursting in Gly^1^-SIFamide application by measuring LPG slow burst duration and cycle period pre-, during, and post-elimination of the pyloric rhythm via hyperpolarization of the two PD neurons. This manipulation sufficiently hyperpolarizes AB, and thus eliminates rhythmic pyloric-timed activity in the pacemaker group (PD, AB, LPG) due to electrical coupling between AB and PD neurons (42, 50). PD hyperpolarization also eliminates LPG pyloric-timed bursting due to electrical coupling, but not LPG gastric mill-timed (slow) bursting in Gly^1^-SIFamide application (Fig. 2) (17). However, the previous study did not determine whether the LPG slow bursting in the absence of rhythmic pyloric input differed from control slow bursting.

**Figure 2.**
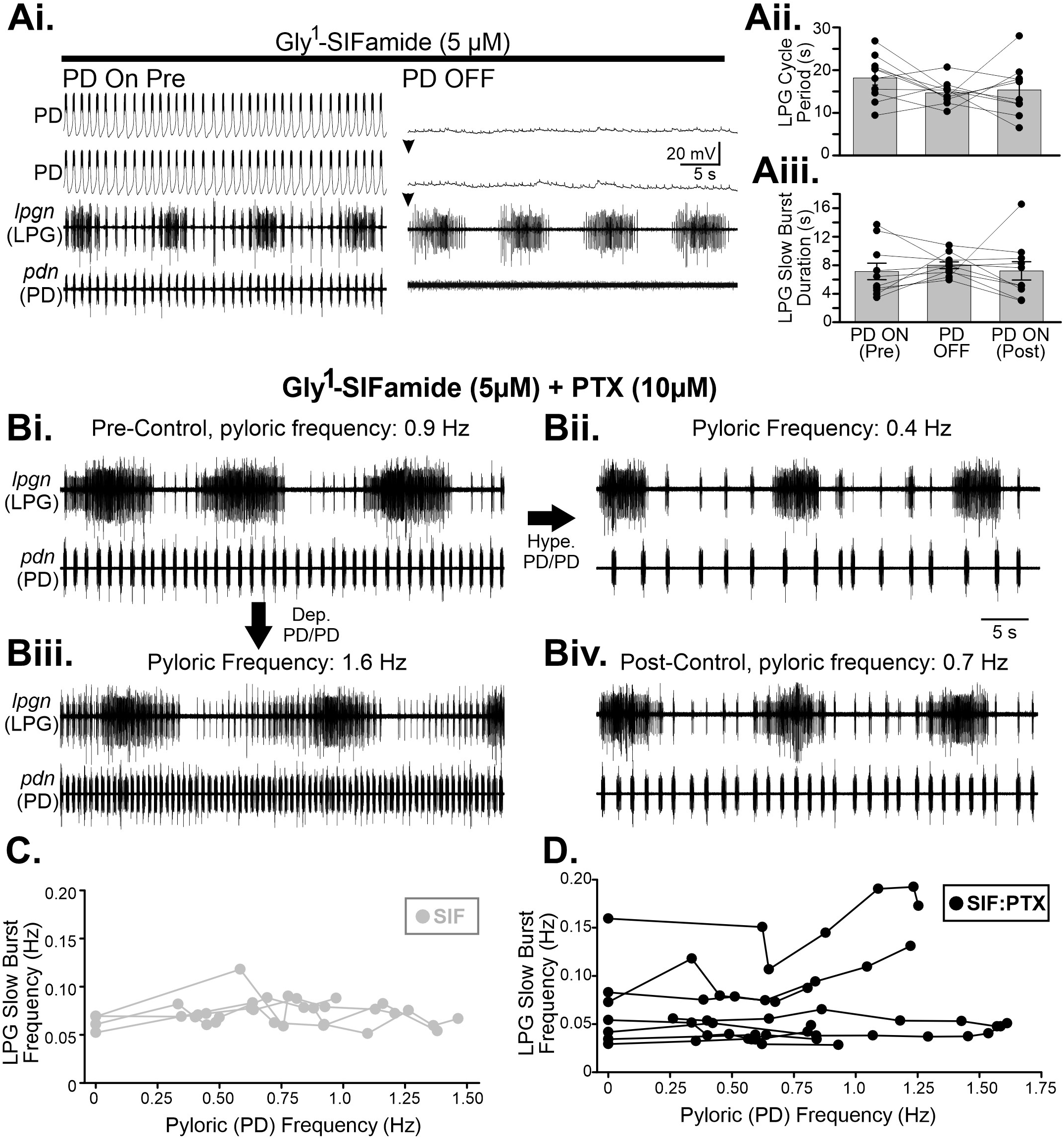
Shutting off the pyloric rhythm or altering pyloric frequency does not regulate LPG gastric mill-timed (slow) bursting during Gly^1^-SIFamide application. ***Ai***, LPG slow bursting occurred during Gly^1^-SIFamide bath application in the control condition (*left*) and when the pyloric rhythm was turned off (*right*) with hyperpolarizing current injection in the two PD neurons (downward arrowheads). LPG slow burst cycle period (Ai) and burst duration (Aii) during PD neuron activity (PD ON-pre/-post) and hyperpolarization (PD OFF). Aii, n = 10, p > 0.05, One-way repeated measures ANOVA; Aiii, n = 10, p > 0.05, Friedman repeated measures ANOVA On Ranks. ***Ci.*** Example recordings of extracellular LPG (*lpgn*) and PD (*pdn*) activity are shown during Gly^1^-Gly^1^-SIFamide plus picrotoxin (PTX) application at baseline pyloric frequency (***Bi***) and during hyperpolarizing current injection into the two PD neurons to slow the pyloric rhythm (***Bii***) or depolarizing current injection in the PDs to increase pyloric rhythm speed (***Biii***), and a post-control (***Biv***) after a series of current injections. LPG slow burst frequency is plotted as a function of pyloric frequency for a series of current injection performed during Gly^1^-SIFamide bath application (***C***) or during Gly^1^-SIFamide plus PTX application (***C***). *C,* r = 0.020, r^2^ = 0.0004, p = 0.909, n = 4, Pearson Correlation; *D*, r = 0.098, r^2^ = 0.009, p = 0.485, n = 8, Pearson Correlation.

Shutting off the pyloric rhythm had no effect on LPG slow burst cycle period (Fig. 2Ai-Aii; PD ON-pre = 18.16 ± 1.66, PD OFF = 14.69 ± 0.88, PD ON-post = 15.36 ± 1.93; p = 0.231, n = 10, One-Way RM ANOVA) or slow burst duration (Fig. 2Aiii,; PD ON-pre = 7.131 ± 1.66, PD OFF 8.00 ± 0.48, PD ON-post 7.22 ± 1.28; p = 0.273, n = 10, Friedman RM ANOVA on Ranks). One possible explanation for the inability of the pyloric network to regulate LPG slow bursting could be that electrical coupling between LPG and PD neurons is rectifying, such that positive current flows more easily from PD to LPG, and negative current flows from LPG to PD (47). Thus, although PD hyperpolarization may not influence LPG slow bursting due to this rectification, the frequency of rhythmic AB/PD depolarizations might regulate LPG slow bursting.

To test for an influence of pyloric frequency on LPG slow bursting, we used steady current injections of hyperpolarizing or depolarizing current (-6 to +2.5 nA) into the two PD neurons to decrease or increase pyloric frequency. Across a range of pyloric frequencies, we found no correlation between pyloric frequency and LPG slow burst frequency (Fig. 2C) (r = 0.020, r^2^ = 0.0004, p = 0.909, n = 4; Pearson correlation), or between pyloric frequency and LPG slow burst duration (r = 0.182, r^2^ = 0.033, p = 0.297, n = 4; Pearson correlation) (data not shown). To ensure that variable synaptic input from other gastric mill neurons (see Results below) was not masking the ability of the pyloric network to regulate LPG slow bursting, we also manipulated pyloric frequency with gastric mill network inputs blocked. Glutamatergic IC, LG, and DG synaptic actions were blocked with picrotoxin (PTX) and the pyloric frequency again regulated with depolarizing and hyperpolarizing current injections during Gly^1^-SIFamide in the presence of PTX. There was some variability such as shorter LPG slow burst duration at a slower frequency in the example shown (Fig. 2Bi vs 2Bii) and a slower LPG slow burst frequency at a higher pyloric frequency (Fig. 2Biii). However, there was no correlation between pyloric frequency and LPG slow burst frequency (Fig. 2D; r = 0.098, r^2^ = 0.009, p = 0.485, n = 8, Pearson Correlation), and no correlation between pyloric frequency and LPG slow burst duration (r = -0.140, r^2^ = 0.020, p = 0.317, n = 8, Pearson Correlation) (data not shown). Thus, the pyloric network was not able to regulate LPG slow bursting activity in Gly^1^-SIFamide.

### Regulation of LPG via gastric mill network neurons IC, DG, and LG

We previously found that, when gastric mill activity was eliminated, the overall percent of LPG slow bursting was not different from when gastric mill activity was present, thus the gastric mill network is not necessary for LPG to generate slow bursting (17). For instance, when gastric mill network neurons LG, IC, and DG are simultaneously hyperpolarized to eliminate their activity (LC/IC/DG OFF), LPG maintains its ability to generate slow bursting (Fig. 3A) (17). However, in the previous study we did not determine whether LPG slow bursting is regulated by the gastric mill network. Thus, we measured LPG slow burst duration and cycle period when LG, IC, and DG gastric mill neuron activity were eliminated via hyperpolarization, individually and collectively (Fig. 3, *re-analyzed dataset from 17*). Our measurements of LPG bursting activity were taken from *lpgn* recordings, in which both copies of LPG are present, making it difficult to assign action potentials to each individual LPG. Thus, we did not quantify LPG number of spikes per burst or firing frequency.

**Figure 3.**
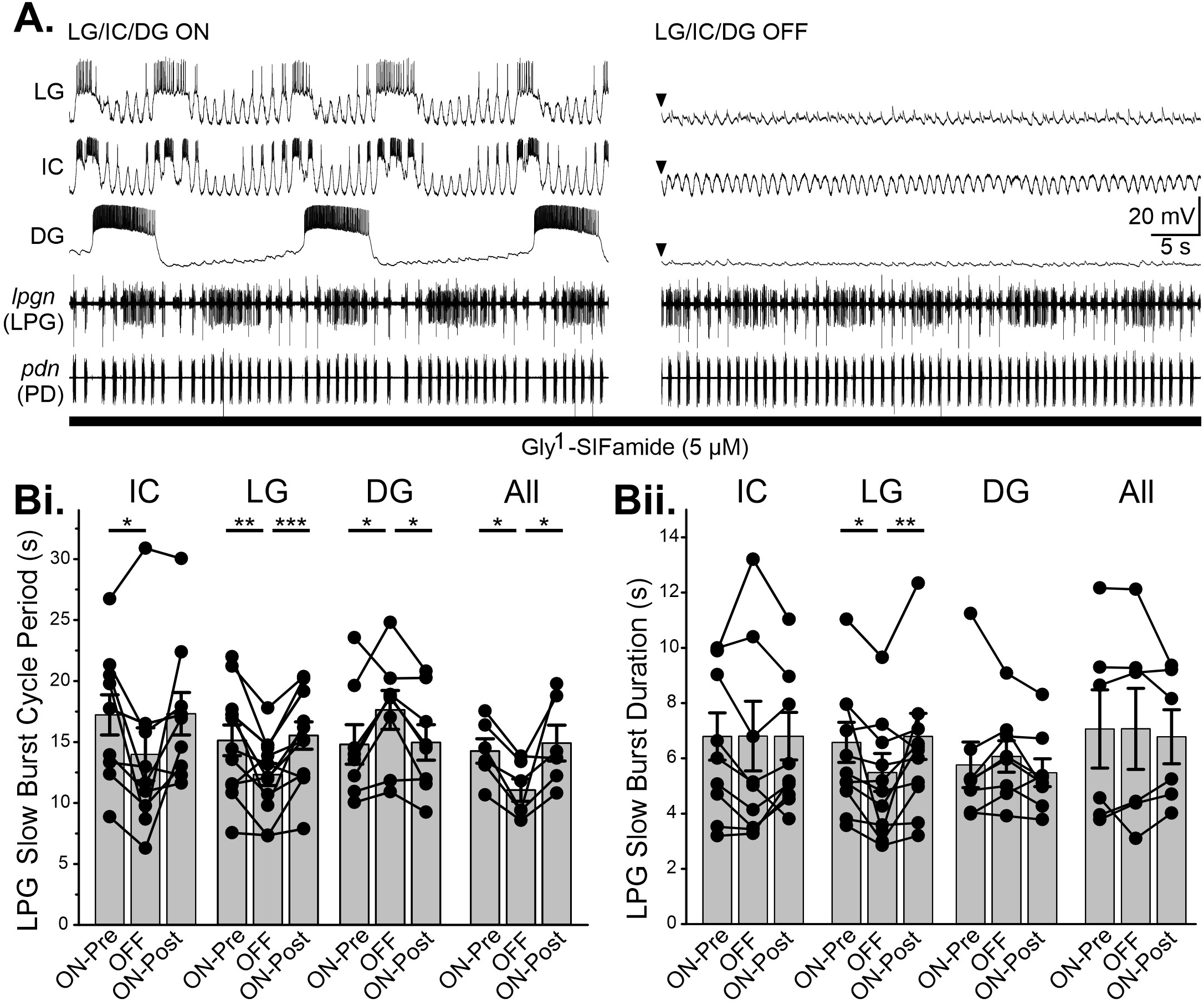
Gastric mill network neurons regulated LPG slow bursting in Gly^1^-SIFamide. ***A***. Intracellular (LG, IC, DG) and extracellular (*lpgn* [LPG], *pdn* [PD]) recordings during Gly^1^-SIFamide application, with LG, IC, and DG neurons active (*left,* LG/IC/DG ON) and during injection of hyperpolarizing current (*right*, LG/IC/DG OFF). LPG generates dual bursting activity during hyperpolarization of LG, IC, and DG. ***Bi, Bii.*** LPG slow burst cycle period (***Bi***) and burst duration (***Bii***) before (ON-pre), during (OFF), and after (ON-post) individual and simultaneous IC, LG, and DG hyperpolarization. n = 10, One-way repeated measures ANOVA, Holm-Sidak *post hoc* analysis, *p < 0.05; **p < 0.01; ***p < 0.001. Data re-analyzed from 17.

Hyperpolarizing gastric mill neurons did influence LPG slow bursting. Specifically, hyperpolarization of IC elicited a decrease in LPG slow burst cycle period but did not affect LPG slow burst duration (Fig. 3Bi, Bii, LPG cycle period: p = 0.029, Table 1; LPG burst duration: p = 1.000, Table 1; n = 10, One-way RM ANOVA). When LG was hyperpolarized, both LPG slow burst cycle period and burst duration were decreased (Fig. 3Bi, Bii, LPG cycle period: p < 0.001, Table 1; LPG burst duration: p = 0.003, Table 1; n = 12, One-way RM ANOVA). DG hyperpolarization elicited an increase in LPG slow burst cycle period but did not affect LPG slow burst duration (Fig. 3Bi, Bii, LPG cycle period: p = 0.008, Table 1; LPG burst duration: p = 0.370, Table 1; n = 8, One-way RM ANOVA). During simultaneous hyperpolarization of LG, IC, and DG neurons (LG/IC/DG OFF), LPG cycle period was shorter, however, LPG slow burst duration was not affected (Fig. 3Bi, Bii, LPG cycle period: p = 0.017, Table 1; LPG burst duration: p = 0.824, Table 1; n = 6, One-way RM ANOVA). Thus, although LPG slow bursting can occur without gastric mill inputs, this slow bursting is regulated by gastric mill network neurons.

**Table 1.** LPG slow burst cycle period and burst duration with/without gastric mill network neuron activity.

| | | Mean ± SEM<br>(In Gly<sup>1</sup>-SIFamide) | | | | *p*-value | | |
| --- | --- | --- | --- | --- | --- | --- | --- | --- |
| Manipulation |  | ON-Pre | OFF | ON-Post | *F/H* | OFF vs | OFF vs | ON-Pre vs |
|  |  |  |  |  |  | ON-Pre | ON-Post | ON-Post |
| IC hype<br>(n = 10,<br>df = 9,2) | Cyc | 17.2 ± 1.7 | 14.0 ± 2.2 | 17.3 ± 1.7 | 4.358 | 0.043 | 0.054 | 0.94 |
|  | Per (s) |  |  |  |  |  |  |  |
| | Burst | 6.8 ± 0.9 | 6.8 ± 1.3 | 6.8 ± 0.9 | 0.0005 | 0.054 | 0.054 | 0.940 |
|  | Dur (s) |  |  |  |  |  |  |  |
| LG hype<br>(n = 12,<br>df = 11,2) | Cyc | 15.1 ± 1.3 | 12.3 ± 0.9 | 15.5 ± 1.1 | 11.72 | 0.002 | <0.001 | 0.587 |
|  | Per (s) |  |  |  |  |  |  |  |
| | Burst | 6.6 ± 0.7 | 5.5 ± 0.7 | 6.8 ± 0.8 | 7.753 | 0.011 | 0.004 | 0.549 |
|  | Dur (s) |  |  |  |  |  |  |  |
| DG hype<br>(n = 8,<br>df = 7,2) | Cyc | 14.8 ± 1.6 | 17.6 ± 1.6 | 15.0 ± 1.5 | 7.012 | 0.015 | 0.015 | 0.849 |
|  | Per (s) |  |  |  |  |  |  |  |
| | Burst | 5.8 ± 0.8 | 6.1 ± 0.6 | 5.5 ± 0.5 | 1.07 | | | |
|  | Dur (s) |  |  |  |  |  |  |  |
| IC/LG/DG<br>hype<br>(n = 6,<br>df = 5,2) | Cyc | 14.3 ± 1.0 | 11.1 ± 0.9 | 14.9 ± 1.5 | 6.281 | 0.04 | 0.023 | 0.591 |
|  | Per (s) |  |  |  |  |  |  |  |
| | Burst | 7.1 ± 1.4 | 7.1 ± 1.5 | 6.8 ± 1.0 | 0.197 | | | |
|  | Dur (s) |  |  |  |  |  |  |  |

During Gly<sup>1</sup>-SIFamide application; ON-Pre, before neuron(s) hyperpolarized; ON-Post, after neuron(s) hyperpolarized; Cyc Per, Cycle Period; Burst Dur, Burst Duration; *Post hoc* not performed when *p* > 0.05. One-way repeated measures ANOVA, Holm-Sidak *post hoc*

### LPG contributions to gastric mill rhythm and pattern generation

Thus far, we found that gastric mill network neurons LG, IC, and DG regulate LPG slow burst activity. However, the role of a switching neuron in a second network when recruited via modulation of intrinsic properties is unknown. Thus, we examined the role of LPG switching into the gastric mill network during Gly^1^-SIFamide application. One way to test this would be to hyperpolarize LPG neurons to eliminate their activity and compare LG, IC, and DG burst parameter measurements between LPG active versus LPG inactive. However, due to rectifying electrical coupling between LPG and PD neurons (47), negative current injections into LPG would hyperpolarize AB/PD and slow or eliminate the pyloric rhythm and thus not test the impact of only altering LPG activity. Instead, we used photoinactivation (see Methods; 44) to selectively eliminate LPG activity without eliminating the pyloric rhythm and compared LG, IC, and DG gastric mill neuron activity in LPG intact versus LPG photoinactivated (LPG Killed) conditions (Fig. 4).

**Figure 4.**
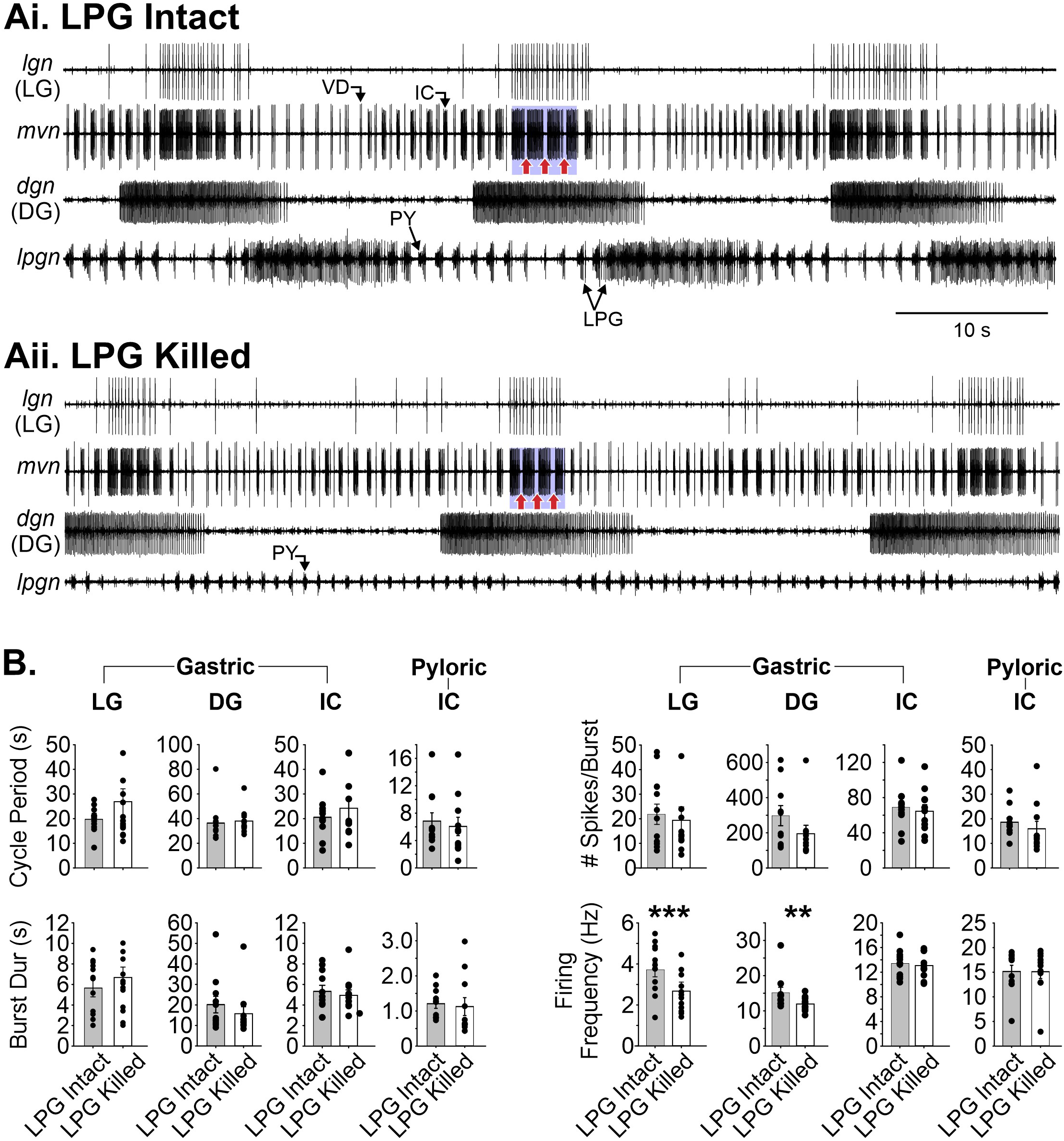
LPG regulated gastric mill network neuron activity but was not necessary for gastric mill rhythm generation. ***Ai, Aii.*** Extracellular recordings of LG (*lgn*), IC (*mvn*), DG (*dgn*), and LPG (*lpgn*) during Gly^1^-SIFamide application with LPG Intact (***Ai***) and LPG photoinactivated (***Aii***, LPG Killed). LG, IC, and DG generated a gastric mill rhythm after LPG was photoinactivated (LPG Killed). Blue boxes indicate IC gastric mill-timed bursting with pyloric timing (red arrows) during LPG Intact and LPG Killed (***Ai*** and ***Aii***, respectively). ***B.*** Cycle period, burst duration, number of spikes per burst, and firing frequency of LG (n = 12), DG (n = 10), and IC (n = 11) during LPG Intact and LPG Killed. The same parameters were measured for IC pyloric-timed bursts (Pyloric-IC). IC gastric mill-timed (see Methods) and pyloric-timed (see Methods) bursts were analyzed separately. Paired t-test, **p < 0.01; ***p < 0.001.

Since Gly^1^-SIFamide modulates LPG to become an intrinsic burster neuron in the gastric mill network during Gly^1^-SIFamide application (5 µM) (17, 23), we first determined whether LPG slow bursting is necessary for generating the Gly^1^-SIFamide-elicited gastric mill rhythm (Fig. 4). When LPG was intact and dual-active (pyloric and gastric mill-timed, *lpgn*), the gastric mill neurons LG (*lgn*), IC (*mvn*), and DG (*dgn*) generated gastric mill-timed bursting activity (Fig. 4Ai). Following photoinactivation of LPG (2 copies, LPG Killed), the LG, IC, and DG neurons maintained their bursting activity, in all preparations tested (Fig. 4Aii, n = 10/10). Thus, LPG slow bursting is not necessary for gastric mill rhythm generation.

Although the gastric mill rhythm still occurred following LPG Killed, we observed a qualitative change in the overall activity of gastric mill network neurons (Fig. 4A). For instance, in the example shown, LG bursts appeared to be shorter in duration with a lower firing frequency (Fig. 4A). Quantifying LG, IC, and DG activity parameters, we found that LG and DG firing frequency was decreased after LPG Killed (Fig. 4B, LG: p = 0.0008, Table 2, n = 12, paired t-test; DG: p = 0.002, Table 3, n = 10, Wilcoxon Signed Rank Test). LPG did not regulate IC firing frequency (Fig. 4B, IC: p = 0.472, Table 2, n = 11, paired t-test), or cycle period, burst duration, or number of spikes per burst for LG, IC, and DG (Fig. 4B; Cycle Period: LG, p = 0.371, n = 12, paired t-test; IC, p = 0.173, n = 11, paired t-test; DG, p = 0.102, n = 11, Wilcoxon Signed Rank Test; Burst Duration: LG, p = 0.928, n = 12, paired t-test; IC, p = 0.438, n = 11, paired t-test; DG, p = 0.123, p = 11, Wilcoxon Signed Rank Test; Number of Spikes per Burst: LG, p = 0.082, n = 12, paired t-test; IC, p = 0.505, n = 11, paired t-test; DG, p = 0.230, p = 10, paired t-test; Table 2).

**Table 2.** LG, DG, and IC burst parameters during LPG Intact and LPG Photoinactivated (Killed) conditions.

| | | Mean ± SEM | | *t/z* | *p-value* |
| --- | --- | --- | --- | --- | --- |
|  |  | LPG Intact | LPG Killed |  |  |
| LG<br>(n = 12,<br>df = 11) | Cyc Per (s) | 19.8 ± 1.5 | 22.3 ± 3.1 | -0.93* | 0.371 |
| | Burst Dur (s) | 6.0 ± 0.7 | 5.8 ± 0.75 | 0.09* | 0.928 |
| | # Spikes/Burst | 23.4 ± 4.1 | 15.9 ± 3.1 | 1.92* | 0.082 |
| | Firing Freq (Hz) | 3.8 ± 0.4 | 2.6 ± 0.3 | 4.61* | 0.0008 |
| DG<br>(n = 10,<br>df = 9) | Cyc Per (s) | 36.4 ± 4.7 | 37.8 ± 3.1 | -1.7** | 0.102 |
| | Burst Dur (s) | 20.2 ± 4.0 | 15.7 ± 3.4 | -1.6** | 0.123 |
| | # Spikes/Burst | 297.7 ± 57.3 | 195.0 ± 48.2 | 1.29* | 0.23 |
| | Firing Freq (Hz) | 15.1 ± 1.6 | 11.9 ± 0.7 | -2.8** | 0.002 |
| IC, slow<br>bursts<br>(n = 11,<br>df = 10) | Cyc Per (s) | 20.6 ± 2.5 | 24.2 ± 3.9 | -1.47* | 0.173 |
| | Burst Dur (s) | 5.3 ± 0.6 | 4.9 ± 0.5 | 0.81* | 0.438 |
| | # Spikes/Burst | 68.9 ± 7.2 | 64.5 ± 7.5 | 0.69* | 0.505 |
| | Firing Freq (Hz) | 13.4 ± 0.7 | 13.0 ± 0.6 | 0.75* | 0.472 |
| IC, fast<br>bursts<br>(n = 11,<br>df = 10) | Cyc Per (s) | 6.83 ± 1.22 | 6.06 ± 1.34 | -1.16** | 0.278 |
| | Burst Dur (s) | 1.21 ± 0.13 | 1.12 ± 0.25 | -0.98** | 0.365 |
| | # Spikes/Burst | 18.61 ± 1.77 | 15.93 ± 3.03 | -1.33** | 0.206 |
| | Firing Freq (Hz) | 15.15 ± 1.26 | 15.10 ± 1.39 | 0.13* | 0.900 |

During Gly<sup>1</sup>-SIFamide application; \*t-statistic (t), paired t-test, \*\*z-statistic (z), Wilcoxon Signed Rank test

**Table 3.** Mean ± SEM percentage of identified gastric mill categories during Gly^1^-SIFamide application with LPG Intact and LPG Killed.

| Coordination Category | Gly<sup>1</sup>-SIFamide (SIF) |  |  | LPG Intact |  | LPG Intact (excl.) |  |
| --- | --- | --- | --- | --- | --- | --- | --- |
|  |  |  |  | (incl. vs excl.) |  | vs. LPG Killed |  |
|  |  |  |  | (n = 9) |  | (n = 10) |  |
|  | LPG Intact | LPG Intact | LPG Killed | t/z | p | t/z | p |
|  | (LPG incl.) | (LPG excl.) |  |  |  |  |  |
| All Coord. | 20.7 ± 3.8 | 34.8 ± 4.6 | 31.0 ± 4.9 | -2.52** | 0.008 | 0.55* | 0.597 |
| All Uncoord. | 25.2 ± 5.1 | 6.6 ± 2.4 | 11.7 ± 4.0 | -5.36* | <0.001 | -0.84** | 0.461 |
| One Uncoord. | 19.6 ± 6.4 | 10.3 ± 5.6 | 10.3 ± 7.1 | -1.48* | 0.177 | -0.0002* | 1.000 |
| One Tonic | 14.9 ± 6.0 | 20.8 ± 6.2 | 6.3 ± 3.3 | -1.83** | 0.125 | 2.18* | 0.057 |
| One Silent | 19.0 ± 4.3 | 23.4 ± 4.0 | 36.1 ± 6.4 | 2.31* | 0.0496 | -1.75* | 0.113 |
| All Tonic | 0 | 0.19 ± 0.19 | 0 | -1.00** | 1.000 | -2.9E-250** | 1.000 |
| All Silent | 0.5 ± 0.3 | 3.9 ± 2.1 | 4.6 ± 2.4 | -1.83** | 0.125 | 0.65** | 0.57 |

\*t-statistic (t), paired t-test, \*\*z-statistic (z), Wilcoxon Signed Rank test. "incl." indicates LPG was included in the category analysis, "excl." indicates that LPG was Intact but was not used to identify categories.

During Gly^1^-SIFamide application, IC gastric-mill timed bursts (Fig. 4A, blue boxes; see Methods for criteria) sometimes consist of multiple bursts that are interrupted in pyloric time (Fig. 4A, red arrows) (17, 36). Thus, in addition to examining LPG effects on IC gastric mill timing, we examined whether IC pyloric-timed activity was regulated by LPG slow bursting and found that pyloric-timed IC bursting was not different after LPG photoinactivation (Fig. 4B, Pyloric-IC; p=0.206 – 0.900, Table 2). Overall, these results indicate that while LPG is not necessary for generating the gastric mill rhythm, it does play a role in regulating some aspects of gastric mill network neuron activity.

### LPG is not necessary for coordinating the Gly^1^-SIFamide gastric mill rhythm

CPG pattern generation involves the relative timing, i.e., coordination of network neuron activity (2, 6, 51–54). Additionally, the strength of neuron activity, including firing frequency, is important for muscle and other targets outside the network, but can also be important for interactions between synaptically connected network neurons, and thus impact coordination (24, 55–58). Since LPG influences gastric mill neuron firing frequency, we next examined whether LPG was necessary for coordinating LG, IC, and DG neurons in Gly^1^-SIFamide.

Typically, in Gly^1^-SIFamide (5 µM), the pattern of the gastric mill rhythm is variable both within and across preparations, posing a challenge to detect changes in network coordination between the intact and LPG killed conditions. Thus, we used a qualitative approach in which we categorized coordination among gastric mill network neurons on a cycle-by-cycle basis, using the LG neuron as the reference neuron (Fig. 5, *see Methods*, adapted from [45]). Briefly, LG, IC, DG, and LPG burst starts and stops were identified across a 1200 s time window, and coordination among three (LG, IC, and DG) or four (LG, IC, DG, and LPG) neurons was determined based on a series of coordination types observed in Gly^1^-SIFamide-elicited gastric mill rhythms. We considered “coordination” to be instances with 1:1 relationships between neuron bursts. For instance, “All Coordinated” indicates a single burst in other network neurons occurring within one LG cycle (Fig. 5A, “All Coordinated”). In addition, “All Coordinated” includes burst onset occurring near the end of the previous LG cycle (less than 10% into the previous cycle) plus burst offset occurring in the current LG cycle; or burst onset occurring in the current LG cycle (at least 10%) and burst offset in the next LG cycle. For instance, in figure 5Bi, the first “All Coordinated” cycle has IC and LPG bursts with onsets and offsets within the LG cycle. In addition, the DG neuron onset in this cycle was greater than 10% and had an offset in the next cycle (Fig. 5Bi, “All Coordinated”). If a neuron was active across a complete LG cycle and extended beyond that LG cycle, the cycle was classified as “One Neuron Tonic” (Fig. 5, “One Neuron Tonic”). DG was the only neuron that exhibited “Tonic” activity. When one or more neurons were silent, that cycle was defined as “One Neuron Silent” or “All Silent”, depending on the number not active in that cycle (Fig. 5, “One Neuron Silent”, “All Silent”). Finally, when IC, DG, and/or LPG neurons had more than one burst per LG cycle, or some other burst pattern that did not meet the above criteria, that cycle was defined as “One Neuron Uncoordinated” or “All Uncoordinated” (Fig. 5). Within each preparation, we found that a single coordination type might only persist for a single cycle at a time. For instance, in the example traces in figure 5Bi, one preparation exhibited cycles categorized as “One Neuron Tonic”, plus an alternation between “All Coordinated” and “One Neuron Silent” cycles (Fig. 5Bi). Three additional preparations provide examples of additional types of coordination that were observed across preparations (Fig. 5Bii). Given the level of variability of the gastric mill rhythm even within each preparation, we quantified the percentages of coordination types across preparations and conditions on a per cycle basis.

**Figure 5.**
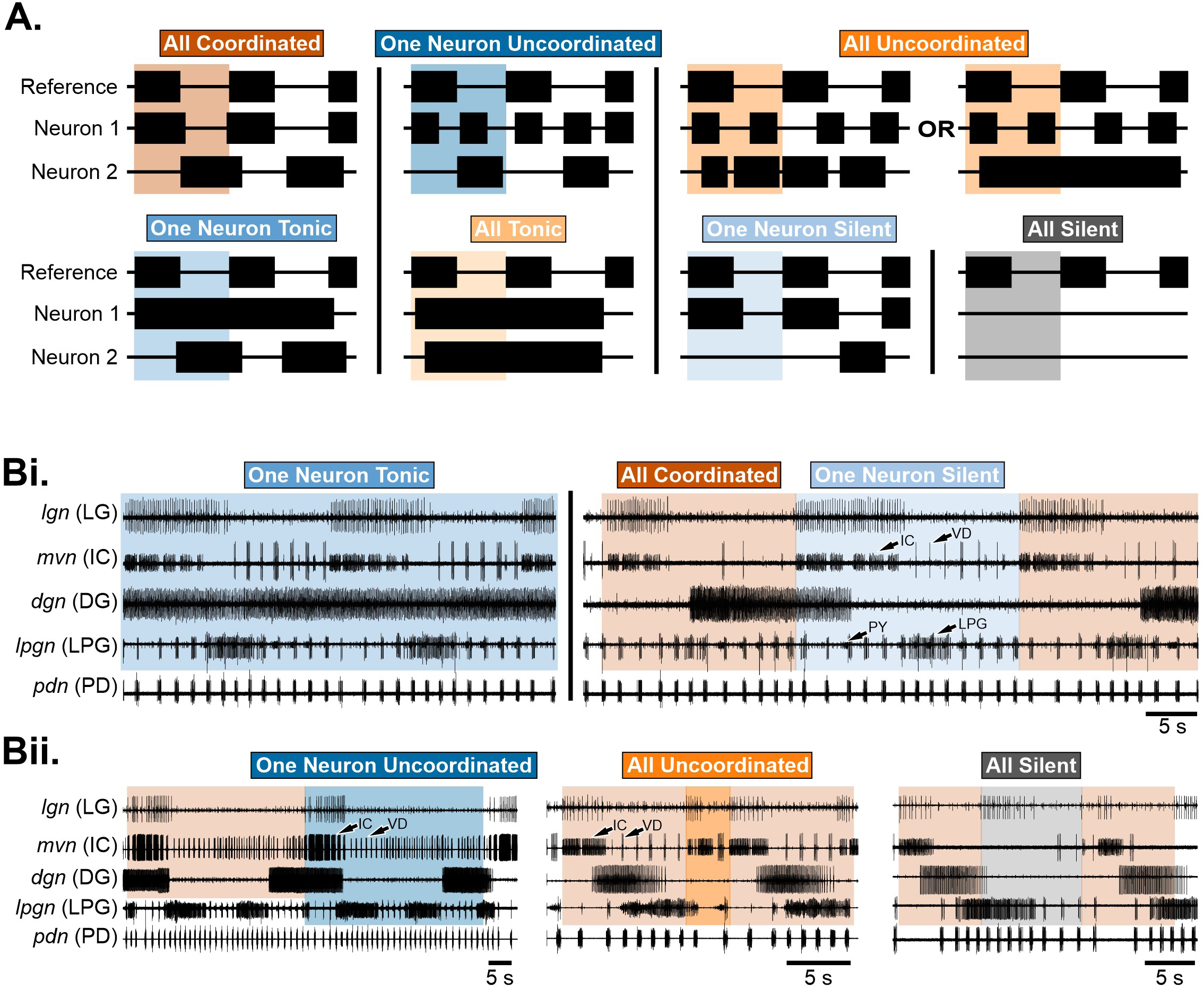
Categorization of gastric mill rhythm coordination during Gly^1^-SIFamide application. ***A.*** Schematics of possible types of coordination between a reference neuron and two network neurons (Neuron 1, Neuron 2). Each colored shaded region indicates one cycle of rhythmic activity with one type of coordination. For example, in “All Coordinated”, Neuron 1 and Neuron 2 each have bursts that begin and end within one cycle of the reference neuron (brown shaded region). In some cases, there can be more than one possibility for a type of coordination. For instance, in the “All Uncoordinated” case, Neurons 1 and 2 may have multiple bursts within one Reference Neuron cycle, or Neuron 1 may have multiple bursts while Neuron 2 fires tonically during the Reference Neuron cycle. See Methods for a complete description of categories. ***Bi-Bii***. Example traces of LG (*lgn*), IC (*mvn*), DG (*dgn*), LPG (*lpgn*), and PD (*pdn*) show different types of coordination between these neurons. ***Bi.*** Different types of coordination occurred within one preparation during the Gly^1^-SIFamide-elicited gastric mill rhythm. In the example shown, coordination was determined among LG, IC, DG, and LPG. At the beginning of the example, there were two cycles of “One Neuron Tonic”, in which DG fired tonically. After some time (vertical solid line), there was one cycle of “All Coordinated” activity, then one cycle of “One Neuron Silent” followed by another cycle of “All Coordinated activity. In the “One Neuron Silent” cycle, the DG neuron was identified as silent, as more than 10% of its activity was attributed to the cycle prior, and thus associated with that cycle. ***Bii.*** Other example traces, each from different preparations, show physiological examples of other coordination categories. *Left*, The coordination category can be different depending on whether LPG is excluded (brown shading, “All Coordinated”) or included (dark blue shading, “One Neuron Uncoordinated”) in the categorization analysis. In the “All Silent” cycle, the LPG neuron was identified as silent, as more than 10% of its activity was attributed to the cycle prior, and thus associated with that cycle.

First, we conducted a set of control experiments to verify that repeated Gly^1^-SIFamide applications have the same effects, and that photoinactivation of two neurons does not alter the gastric mill network response to Gly^1^-SIFamide. In these control experiments we categorized gastric mill rhythm coordination in conditions where (1) two consecutive Gly^1^-SIFamide applications occurred with no neurons photoinactivated and (2) two gastric mill (GM) neurons were photoinactivated before a second Gly^1^-SIFamide application. GM neurons are not excited by Gly^1^-SIFamide (36) and provide an assay for any generalized effects of photoinactivation on the STG. Consecutive Gly^1^-SIFamide applications and photoinactivation of two GM neurons did not affect gastric mill rhythm coordination (Supplemental Table S1-2, Supplemental Fig. S1). These results gave us confidence that we could use multiple applications of Gly^1^-SIFamide before versus after photoinactivating both copies of the LPG neurons and assess whether there were effects on coordination due to the selective elimination of LPG.

Identified gastric mill rhythm patterns from each preparation were accumulated and the mean percent of cycles per category was calculated and represented in a stacked bar graph, where each colored bar represents one category, and each collection of bars indicates one experimental condition analyzed (Fig. 6). In the “SIF:LPG Intact (LPG Included)” condition, IC, LPG, and DG neuron activity was examined relative to LG activity to determine network coordination. In this control condition, 20.71 ± 3.77% of cycles were “All Coordinated”, 19.56 ± 6.42% were “One Neuron Uncoordinated”, 14.91 ± 6.01% were “One Neuron Tonic”, 19.01 ± 4.34% cycles were “One Neuron Silent”, 25.24 ± 5.08% cycles were “All Uncoordinated”, 0% were “All Tonic”, and 0.48 ± 0.26% were “All Silent” (Fig. 6A, Bi). While there are some differences between bath applied and neuronally released neuropeptide (36), it did not appear that variability in the gastric mill rhythm was due to bath application of Gly^1^-SIFamide. We tested this by tonically stimulating the modulatory projection neuron MCN5 (30 Hz stimulation, 200 s time window), and found that there were 24.87 ± 14.71% “All Coordinated” cycles, 4.0 ± 4.0% “One Neuron Uncoordinated” cycles, 3.08 ± 3.08% “One Neuron Tonic” cycles, 21.73 ± 11.20% “One Neuron Silent” cycles, 42.99 ± 9.16% “One Neuron Uncoordinated” cycles, 0% “All Tonic” cycles, and 3.33 ± 3.33% “All Silent” cycles (data not shown). We did not perform a statistical analysis on bath applied versus neuronal release of Gly^1^-SIFamide due to differences in analysis time windows (1200 s vs. 200 s). However, it appears qualitatively that the extent of gastric mill neuron coordination is similar in bath-applied Gly^1^-SIFamide compared to neuronal release from MCN5.

**Figure 6.**
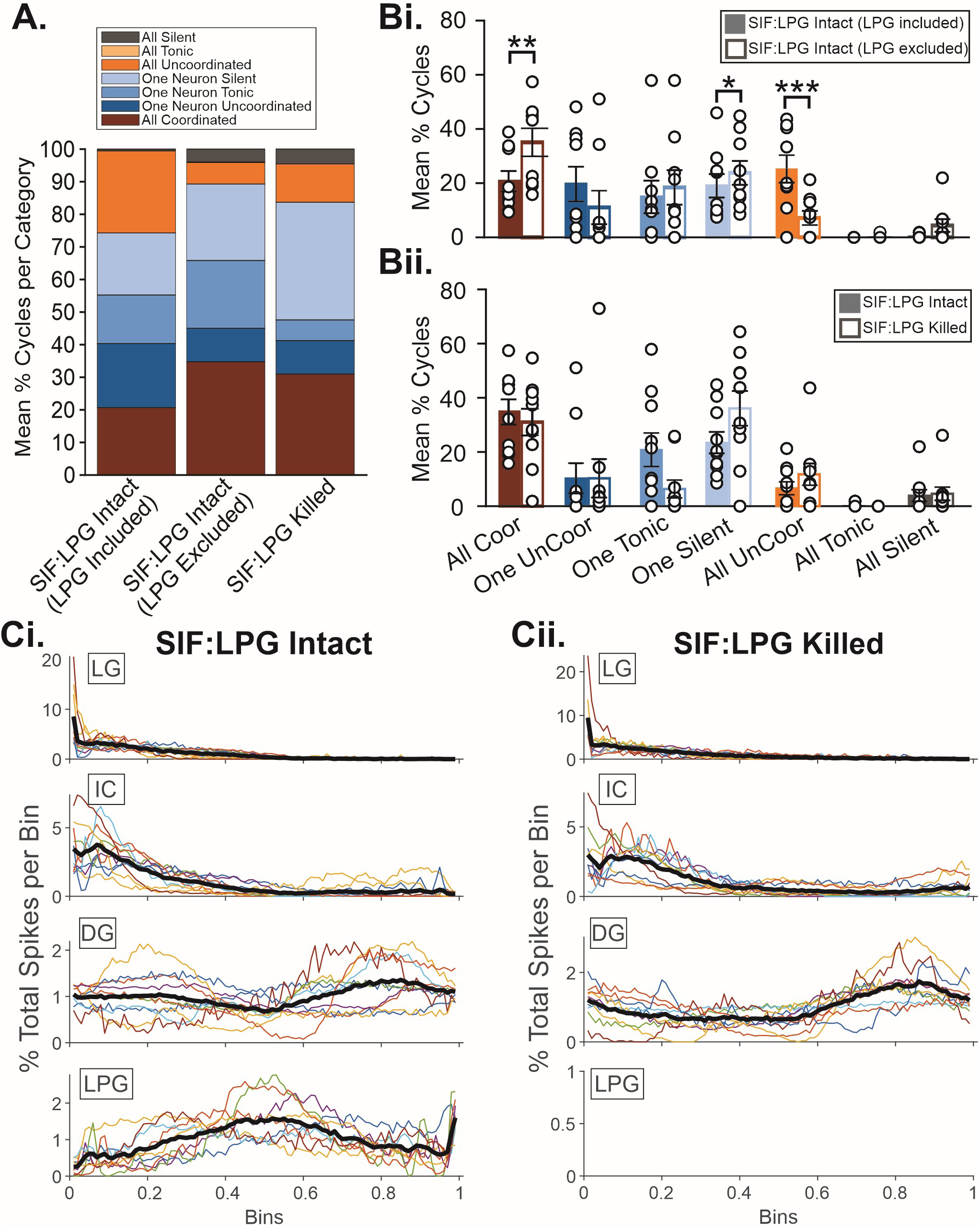
LPG is not necessary for coordinating gastric mill network neuron activity during Gly^1^-SIFamide application. ***A.*** Stacked bar graph showing the mean percentage of coordination categories for LPG Intact with LPG included in the analysis (LPG Included, n = 9), LPG Intact with LPG excluded from the analysis (LPG Excluded, n = 9), and LPG Killed (n = 10). ***Bi-Bii.*** Mean percent of gastric mill coordination types for LPG Intact with LPG (LPG Included, n = 9) versus LPG Intact (LPG Excluded, n = 9) (***Bi***) and LPG Intact (LPG Excluded, n = 10) versus LPG Killed (n = 10) (***Bii***). White data points show the percentage of each coordination category from each preparation. Bars indicate mean ± SEM across preparations. ***Ci, Cii.*** Spike phase analysis of the percent total spikes per bin (100 bins) for LG, IC, DG, and LPG neurons in Gly^1^-SIFamide (SIF) LPG Intact (***Ci***) and SIF LPG Killed (***Cii***) conditions. LG is used as a reference neuron. Colored lines for each neuron indicate individual preparations, while the thick black line indicates the mean percent total of spikes per bin across preparations.

When LPG was excluded [SIF:LPG Intact (LPG Excluded)] from the categorization analysis in the SIF:LPG Intact condition, the percentage of cycles for “All Coordinated” and “One Neuron Silent” was higher and “All Uncoordinated” was lower, while there was no difference for each of the other categories (Fig. 6Bi, Table 3, All Coordinated: p = 0.008, paired t-test; One Neuron Silent: p = 0.0496, paired t-test; All Uncoordinated: p < 0.001, Wilcoxon signed rank test; n = 9). These differences in mean percentage of cycles per category are likely due to a fourth component (LPG) included in pattern identification in the SIF:LPG Intact (LPG Included) measurements. For instance, in two consecutive cycles in figure 5Bii (leftmost traces), the IC and DG neurons are “All Coordinated” with LG, however, with the addition of LPG, the overall coordination indicates that the two cycles are “One Neuron Uncoordinated”. While the inclusion of LPG in the categorization analysis provides more information about the nature of coordination among the gastric mill neurons and LPG slow bursting, it does not address our question of whether LPG plays an important role in mediating coordination among LG, IC, and DG neurons. Thus, to address this question, we chose to identify types of network coordination by including only LG, IC and DG neurons to maintain consistency between SIF:LPG Intact and SIF:LPG Killed conditions.

When LPG was photoinactivated, we found that the mean percentage of LG cycles that were “All Coordinated” was unchanged between SIF:LPG Intact and SIF:LPG photoinactivated (Killed) conditions (Fig. 6A, Bii; Table 3, SIF:LPG Intact: 34.78 ± 4.63%, SIF:LPG Killed: 31.01 ± 4.92%; p = 0.597, n = 10, paired t-test). This suggests that LPG slow bursting is not necessary for coordinating the gastric mill rhythm in Gly^1^-SIFamide. Furthermore, although gastric mill rhythm coordination patterns were variable from cycle-to-cycle (Fig. 5Bi), we observed that the variability in coordination types was similar between SIF:LPG Intact and SIF:LPG Killed conditions (Fig. 6Bii, Coeffficient of variation [CV] for SIF:LPG Intact vs SIF:LPG Killed: All Coordinated, 0.42 vs 0.50; All Uncoordinated, 1.15 vs 1.08; One Uncoordinated, 1.72 vs 2.19; One Tonic, 0.94 vs 1.64; One Silent, 0.53 vs 0.56; All tonic, 3.16 vs N/A; All Silent, 1.72 vs 1.69). Thus, the variability of gastric mill coordination was not due to LPG slow bursting.

While our category approach indicated that there was no difference in the percentage of 1:1:1 bursting activity of LG, IC, and DG neurons with or without LPG, it did not allow us to address the specific pattern and whether there was a change in the relative timing, i.e., phase relationships of the neurons. We wanted to describe the specific relative timing of neuronal activity, for instance, to determine whether DG and LG are coincident or out of phase with each other, regardless of whether there is 1:1 coordination of all neurons in each cycle. To do this, we quantified spiking activity of all neurons, across all cycles, regardless of the type of coordination, again setting LG as the reference neuron. This allowed us to visualize all neuronal activity without limiting the analysis to ∼30% of the cycles that were fully coordinated. To determine whether LPG was involved in mediating the relative timing among LG, IC, and DG network neurons we compared the pattern of spiking with SIF:LPG intact versus SIF:LPG killed (Fig. 6C). Each cycle began with the onset of an LG burst and ended at the start of the subsequent LG burst and was divided into 100 bins. In the control condition, LG spiking extended for ∼40% of the cycle which largely overlapped with spiking in the IC neuron (Fig. 6Ci). DG spiking occurred throughout the cycle with a tendency to have fewer spikes per bin mid-cycle across preparations (colored lines) and also evident in the average activity (thick black line, n = 10), which overlapped with the highest degree of spiking in LPG (LPG slow bursts). In the SIF:LPG Killed condition (Fig. 6Cii), the pattern of activity appeared similar, with LG and IC coactive for the first ∼40% of the cycle, and DG activity was spread across the phase, although it appeared biased toward the end of the cycle. A few preparations which had more DG activity early in the phase in the control condition, did not have such high activity after LPG was killed, but overall there was not a dramatic shift in the pattern, indicating that LPG was not necessary for coordinating the other gastric mill neurons.

### LPG is capable of coordinating gastric mill network neurons

Thus far, we examined the role of LPG in mediating coordination among gastric mill neurons by removing LPG synaptic effects via photoinactivation. Although LPG was not necessary for gastric mill coordination (Fig. 6), we did find that LPG regulated LG and DG firing frequency (Fig. 4), suggesting some role for LPG in determining gastric mill network activity in Gly^1^-SIFamide. In CPG networks, there can be degeneracy at the cellular and synaptic levels, which can provide a “safety net” for networks in the event of injury or other perturbation (91). It was possible that there is synaptic degeneracy among gastric mill neurons in Gly^1^-SIFamide, such that eliminating LPG via photoinactivation does not disrupt coordination due to the other synapses present among LG, IC, and DG neurons. Thus, LPG photoinactivation alone may not enable us to conclude whether LPG can play an active role in the gastric mill network. In the crab STG, the LPG neuron is cholinergic, while LG, IC, and DG neurons are glutamatergic. This distinction in neurotransmitters enabled us to test whether cholinergic LPG synapses can coordinate gastric mill neurons (Fig. 7A).

**Figure 7.**
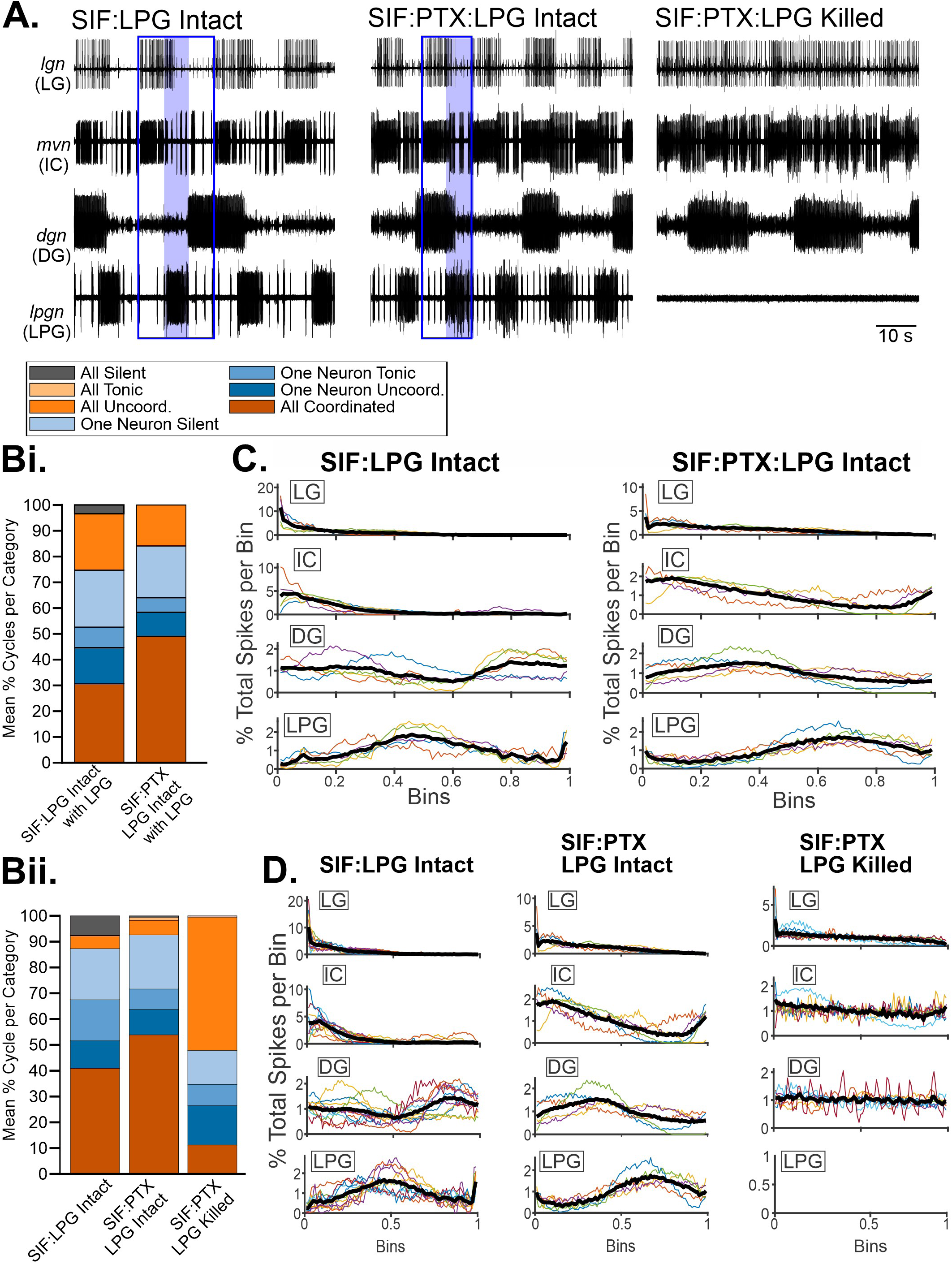
LPG has an active role in mediating coordination among gastric mill network neurons. ***A.*** Example trace recordings of LPG (*lpgn*), LG (*lgn*), IC (*mvn*), and DG (*dgn*) during application of Gly^1^-SIFamide with LPG Intact (SIF:LPG Intact, *left*), SIF plus picrotoxin (PTX)with LPG Intact (SIF:PTX:LPG Intact, *middle*), and SIF:PTX:LPG Killed (*right*). During SIF:LPG Intact, there is a triphasic gastric mill rhythm (blue outline), with LPG gastric mill-timed bursting occurring in the second phase (blue filled box). During SIF:PTX:LPG Intact, glutamatergic inhibitory synapses between LG, IC, and DG are blocked by PTX, and cholinergic LPG synapses are active. In this condition, LPG bursting (blue filled box) alternated with LG, IC, and DG bursting (blue outline). In SIF:PTX:LPG Killed, the gastric mill rhythm was uncoordinated, with tonic LG activity, irregular IC bursting, and DG bursting activity. ***Bi, Bii,*** The mean percentage of coordination categories are plotted as stacked bars for Gly^1^-SIFamide (SIF) LPG Intact (LPG included) versus SIF:PTX LPG Intact (LPG included) (n = 5) (***Bi***) and for SIF LPG Intact (LPG excluded, n = 11) versus SIF:PTX LPG Intact (LPG excluded, n = 5) versus SIF:PTX LPG Killed (n = 8) (***Bii***). ***C.*** Spike phase analysis of the percent total of spikes per bin (100 bins) for LG, IC, DG, and LPG neurons in SIF:LPG Intact (*left*) and SIF:PTX:LPG Intact (*right*) conditions. ***D.*** Spike phase analysis of the percent total of spikes per bin (100 bins) for LG, IC, DG, and LPG neurons in SIF:LPG Intact (*left*), SIF:PTX:LPG Intact (*middle*), and SIF:PTX:LPG Killed (*left*) conditions. In ***C*** and ***D***, LG is used as a reference neuron. Colored lines for each neuron indicate individual preparations, while the thick black line indicates the mean percent total of spikes per bin across preparations.

We found that LPG could coordinate gastric mill neurons on its own, but in a different pattern. For example, in an example cycle in the control Gly^1^-SIFamide condition (SIF:LPG Intact, Fig. 7A, left, blue outline), LG and IC are coactive, LPG is active near the middle of the cycle (blue box) and DG activity begins after LPG. However, when inhibitory glutamatergic synapses were blocked with picrotoxin (PTX, 10 µM) during Gly^1^-SIFamide application (SIF:PTX:LPG Intact), LG, IC, and DG neuron bursting overlapped at the beginning of cycle (e.g., blue outline) and alternated with LPG bursting (Fig. 7A middle, blue box, n = 4/5). Across five experiments in which we applied Gly^1^-SIFamide, followed by Gly^1^-SIFamide + PTX (SIF:PTX:LPG Intact), we applied our categorical analysis to all four neurons (LG, IC, LPG, and DG) and found that the mean percentage of “All Coordinated” activity was not different in SIF:PTX LPG Intact compared to SIF:LPG Intact (Fig. 7Bi, Table 4, p = 0.148, n = 5, paired t-test), Furthermore, there were no differences between SIF:PTX LPG Intact and SIF:LPG Intact for any of the other categories (Table 4). However, although the number of coordinated cycles was not different, there was a qualitative change in the pattern of neuronal activity (Fig. 7A). To quantify this, we again analyzed neuronal spiking (100 bins) across each cycle using LG as the reference neuron. In SIF:PTX:LPG Intact, there was a shift compared to SIF:LPG Intact, with DG now largely coactive with LG and IC, and LPG primarily active at the end of the cycle (Fig. 7C, right). Overall, the pattern shifted from triphasic (IC/LG, LPG, DG) to biphasic (LG/IC/DG, LPG), and indicated that LPG can be sufficient to coordinate gastric mill neurons, even though it is not necessary for the baseline pattern generation.

**Table 4.** Mean ± SEM percentage of identified gastric mill categories during Gly^1^-SIFamide (SIF) application versus Gly^1^-SIFamide plus PTX (SIF:PTX) with LPG Intact.

| Coordination<br>Category | LPG Included (n = 5) |  | *t/z* | *p* |
| --- | --- | --- | --- | --- |
|  | SIF | SIF:PTX |  |  |
| All Coord. | 30.38 ± 4.53 | 49.17 ± 11.98 | -1.792* | 0.148 |
| All Uncoord. | 21.58 ± 4.70 | 15.92 ± 5.34 | 1.335* | 0.253 |
| One Uncoord. | 14.02 ± 8.32 | 9.40 ± 3.42 | 0.476* | 0.659 |
| One Tonic | 7.93 ± 2.59 | 5.82 ± 2.79 | 0.450* | 0.676 |
| One Silent | 22.11 ± 7.38 | 19.69 ± 6.67 | 0.276* | 0.796 |
| All Tonic | 0 | 0 | 2.91E-250** | 1.000 |
| All Silent | 3.97 ± 2.52 | 0 | -1.342** | 0.50 |

Coordination categories identified in Gly<sup>1</sup>-SIFamide (SIF) versus SIF:PTX with LPG included in analysis, \*t-statistic (t), paired t-test, \*\*z-statistic (z), Wilcoxon Signed Rank test

Given the prevalence of electrical coupling in the STG as in other networks (49), it remained possible that in the absence of glutamatergic inhibition among the LG, IC, and DG neurons, electrical coupling coordinated their activity. To determine whether it was electrical coupling, or the LPG cholinergic inhibition that was responsible for the coordination in SIF:PTX:LPG Intact, we combined SIF:PTX application with LPG neuron photoinactivation. In this condition, LG, IC, and DG synapses are blocked via PTX, and LPG activity was selectively eliminated, thus eliminating all intra-network chemical synapses among these gastric mill neurons (35). In this condition, although there was some bursting, particularly in the DG neuron (Fig. 7A, right, *dgn*; n = 6/8), there was no obvious coordination (Fig. 7A, right, SIF:PTX LPG Killed).

To compare the LPG Killed condition with other conditions, we did not include LPG in the categorical analysis and used a larger data set of three conditions (SIF:LPG Intact, n=10; SIF:PTX:LPG Intact. n=5; SIF:PTX:LPG Killed, n=8). We found that the “All Coordinated” category decreased, while “All Uncoordinated” increased (Fig. 7Bii, Table 5; SIF:LPG Intact [n = 5] vs. SIF:PTX LPG Intact [n = 5] vs. SIF:PTX:LPG Killed [n = 8], All Coordinated: p = 0.004, One-way ANOVA; All Uncoordinated: p = 0.04, Kruskal-Wallis one-way ANOVA on ranks). All other categories were unchanged across conditions (Fig. 7Bii; Table 5). Therefore, the overlapping activity observed between LG, IC, and DG in SIF:PTX was not due to electrical coupling among these neurons, but due to synaptic input from LPG. The remaining small percentage of “All Coordinated” cycles in the SIF:PTX:LPG Killed condition suggested a lack of timing relationship among the neurons. This was further evident when examining spiking activity across all cycles in the SIF:PTX:LPG Killed conditions (Fig. 7D, right). Although some bursting activity was evident in some preparations (Fig. 7A, right; Fig. 7D, right, jagged individual experiment lines, IC and DG), overall the activity of the three remaining gastric mill neurons was relatively uniformly distributed across LG “cycles”. Note, we used standard interspike intervals to objectively identify LG bursts (see Methods). For some experiments in the SIF:PTX:LPG Killed condition, this resulted in very few LG bursts due to almost tonic LG activity (number of LG bursts: 8 to 124). Thus, although not necessary for rhythm generation or network coordination, LPG gastric mill-timed bursting is sufficient to coordinate gastric mill network neurons in the Gly^1^-SIFamide modulatory state in the absence of other intra-network chemical synapses.

**Table 5.**
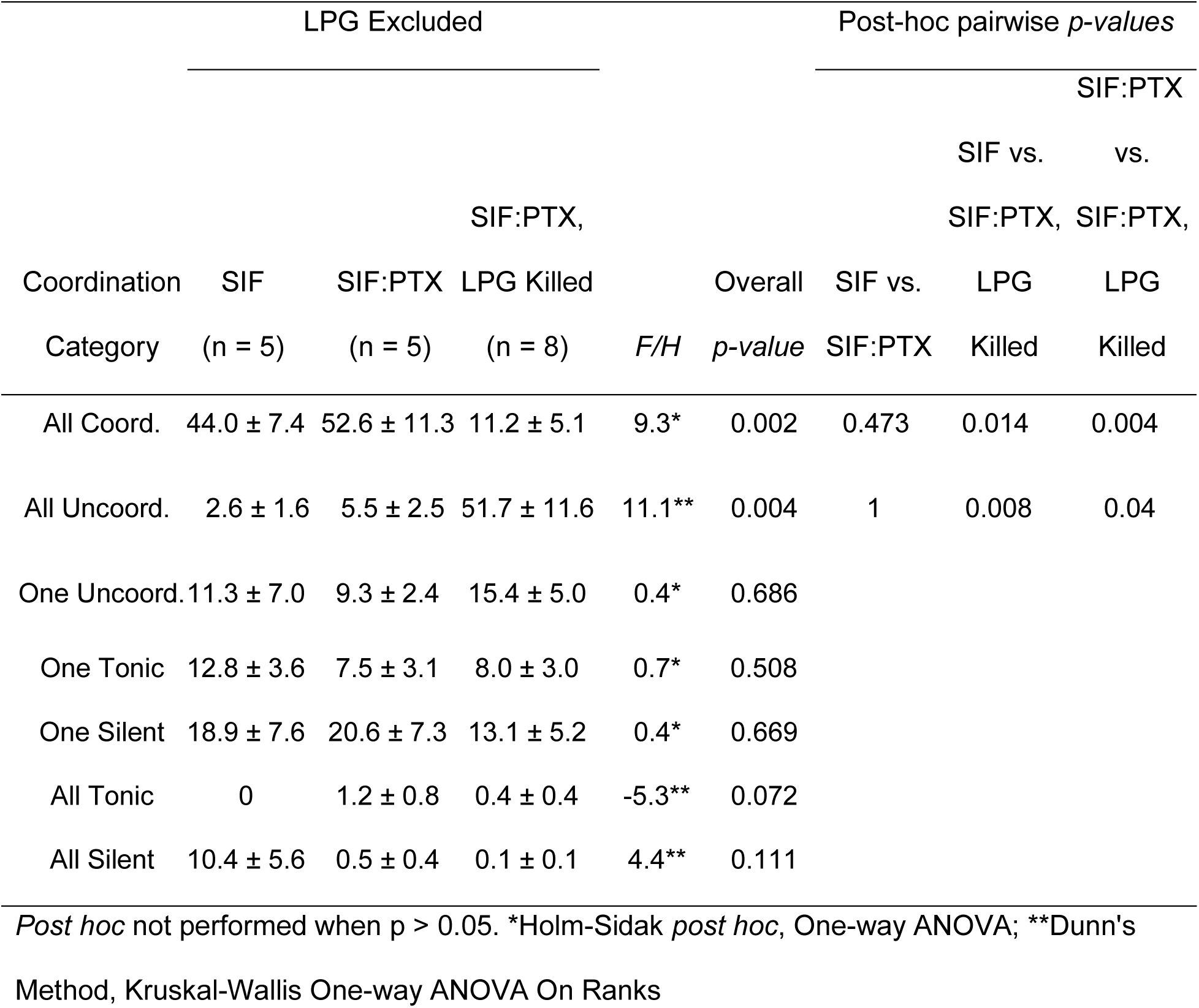
Mean ± SEM percentage of identified gastric mill categories during Gly^1^-SIFamide (SIF) application versus Gly^1^-SIFamide plus PTX with LPG Intact (SIF:PTX) versus SIF:PTX with LPG Killed.

## Discussion

To understand the implications of neuronal switching, it is important to determine the bidirectional interactions between switching neurons and each of their networks. In the present study, we found that the slow gastric mill (∼0.1 Hz), but not the faster pyloric (∼1 Hz), network regulated LPG slow bursting. Additionally, we tested whether LPG regulates the output of its second network, when in its dual-network state. We found that LPG is not necessary for gastric mill rhythm generation, but does regulate aspects of network neuron activity and has the ability to coordinate the entire network. These findings provide novel insight into the roles of a switching neuron which generates dual-network activity via a combination of electrical synapse (pyloric) and intrinsic (gastric mill) mechanisms.

### Switching neuron intrinsically-generated activity is not regulated by its “home” network

The inability of physiological changes in the pyloric network to regulate LPG gastric mill-timed bursting is surprising because LPG gastric mill-timed bursting is intrinsically generated and voltage-sensitive (17). Although the electrical coupling between LPG and the AB/PD pacemaker group is rectifying, it is an incomplete rectification. Specifically, positive current flows preferentially, but not exclusively, from PD to LPG, and similarly for negative current from LPG to PD (47). Thus, we expected that a depolarizing or hyperpolarizing shift in the voltage of the PD neurons, sufficient to alter the voltage-dependent intrinsic oscillations of the pyloric pacemaker, would affect LPG slow bursting, at the very least when PD neurons were depolarized. One possible explanation for a lack of regulation from AB/PD fast bursting to LPG slow bursting is a decrease in electrical coupling strength between these neurons in the Gly^1^-SIFamide modulatory state. However, this possibility is unlikely, as LPG can regulate AB/PD activity during its slow bursts ([36], B Gnanabharathi, S-RH Fahoum, DM Blitz, unpublished observations). An alternative possibility for the lack of pyloric regulation of LPG slow bursting is the time-dependence of the intrinsic currents underlying LPG slow bursting. For instance, in a gastric mill rhythm elicited by the modulatory projection neuron MCN1, rhythmic release of its neuropeptide, CabTRP-Ia causes rhythmic waxing and waning of a modulator-activated inward current (IMI) necessary for gastric mill neuron bursting. A similar gastric mill rhythm version is produced by bath-application of the neuropeptide CabPK, but in this case a time-(and calcium-) dependent current is also necessary for gastric mill neuron rhythmic activity (33, 42, 59, 60). However, in both gastric mill rhythms, the pyloric rhythm does regulate the gastric mill rhythm. A difference between the rhythmic oscillations in gastric mill neurons in the preceding examples and the LPG slow bursting, is that the LPG slow bursts are intrinsically generated and do not require input from other gastric mill neurons (17). In the MCN1- and CabPK-elicited rhythms, in addition to the mentioned intrinsic currents, reciprocal inhibition between LG and Int1 is necessary for their rhythmic activity (33, 61). Our findings suggest that the intrinsic currents underlying the LPG slow bursting (23) may overwhelm the current through the electrical synapses, even in the preferential direction of depolarizing current flow from PD to LPG. However, the lack of control by the pyloric network may be due to more than just the amplitude of the currents, as the interactions between electrical coupling and intrinsic currents can be complex and generate unexpected cellular and circuit level effects (49, 62–64).

### Switching neuron intrinsically generated activity is regulated by its second network

Unlike turning off the pyloric rhythm, eliminating the activity of individual, or all three gastric mill network neurons (LG, DG, IC) altered LPG neuron gastric mill-timed (slow) bursting. These gastric mill neurons altered LPG burst cycle period and/or burst duration despite LPG slow bursting being generated via intrinsic currents (17, 23). In baseline conditions IC, LG, and DG do not have functional synapses onto LPG. Thus, Gly^1^-SIFamide likely modulates intra-network synapses between LPG and the other gastric mill neurons (Fahoum and Blitz, unpublished observations). Intrinsic bursting that is further shaped by synaptic input from the network is a common motif in CPG and other rhythmic neural networks (65–68). For example, the pyloric pacemaker ensemble does not require any synaptic input to generate the rhythm, but cycle period variability is decreased by rhythmic feedback from the LP neuron (58, 69).

### The role of a switching neuron in rhythm and pattern generation in a second network

Initially, it was thought that switching neurons recruited into a second network simply act as follower neurons to carry out patterns of activity determined by the synapses that recruit them (10, 12, 14). However, the role of a switching neuron recruited into a second network via modulation of intrinsic properties had not been examined. One possible role for switching neuron activity in a second network is rhythm generation, to drive CPG rhythmicity and set its frequency (6, 70–74). Because LPG generates endogenous gastric mill-timed bursting during Gly^1^-SIFamide application (17), it was possible that LPG was integral for rhythm generation. However, we found that eliminating LPG activity did not eliminate pyloric or gastric mill rhythmic activity, demonstrating that LPG activity, including its intrinsically generated slow bursting, is not necessary for rhythm generation in either network.

Another possible role for a switching neuron is pattern generation, where neurons determine the activity levels and relative timing among network neurons (71, 75, 76). Here, we found that LPG regulates LG and DG firing frequency. These neurons are members of the gastric mill CPG and motor neurons controlling the movements of the lateral teeth and medial tooth, respectively. The LG and DG firing frequency is thus relevant for the efficacy of synaptic transmission centrally within the CPG, and at the neuromuscular junction (77–79). As a consequence, the impact of LPG on DG and LG activity level can alter behavior directly via changes in muscle contractions, and potentially indirectly through changes in CPG targets of LG and DG.

We found that LPG was not necessary to maintain coordination among gastric mill neurons and there was little difference in the timing relationships in the absence of LPG. Given the variability of the Gly^1^-SIFamide gastric mill rhythm, our analysis might not have been sensitive enough to detect changes in coordination in the absence of LPG, or there might be degeneracy such that other synapses contribute the same function as the LPG neurons (80, 81). However, when other chemical synapses among gastric mill neurons were blocked with picrotoxin, LPG fully coordinated the activity of the other gastric mill network neurons, but with different timing relationships. The DG neuron was both the most variable, and the most different when LPG was the sole contributor to network coordination. The variability in DG activity may reflect the absence of sensory feedback, such as muscle properioceptor feedback (82–85), although DG is more regular during other in vitro gastric mill rhythms (33, 86, 87). Unlike DG, the LG and IC neurons were were consistently out of phase with LPG across conditions. The prolonged IC bursting occurring in the Gly^1^-SIFamide gastric mill rhythm is a hallmark of this rhythm, which does not occur in multiple other gastric mill rhythm versions (25, 61, 86). The ability of LPG to regulate the timing of IC as well as LG emphasizes the important role that LPG is capable of playing, despite it being a temporary visitor from the pyloric network. Thus, modulator-elicited intrinsic LPG slow bursting enables LPG to generate activity at the gastric mill rhythm frequency, plus contribute to patterning the network output and thereby determine the behavioral version.

### Degeneracy as a mechanism for compensation among network neurons

Degeneracy and variability are important themes in motor systems, as they promote network stability and flexibility, respectively (88–91). Various cellular-level mechanisms contribute to degeneracy and variability, including dendritic spine remodeling, perineural net formation, and regulation of axon myelination (91–93). Degeneracy also includes neuromodulatory actions, in which distinct modulators converge on the same ionic current to stabilize neuron bursting activity (5, 94–96). Furthermore, degeneracy of ion channels or synapses contributes to different mechanisms for the same motor pattern under different modulatory states (60, 61, 81). In the Gly^1^-SIFamide modulatory state, partial synaptic degeneracy among LG, IC, LPG, and DG neurons may be important for enabling the gastric mill network to carry out multiple coordination patterns. We do not know if there are distinct behavioral states, or sensory stimuli that might bias the network toward one coordination pattern or another, or whether both types might occur within the same modulatory state. Variability in phase relationships can be periodic, such as the regular switches between peristaltic and near synchronous heart tube contractions in the medicinal leech neurogenic heartbeat system (97).

Degeneracy and variability also serve an important role in rehabilitation in the event of injury or disease. At the network level, degeneracy and variability must work hand-in-hand, such that degeneracy stabilizes and maintains behavior post-injury, while variability promotes discovering new pathways to carry out behaviors (91, 98, 99). For instance, in brain injury and rodent stroke models, dendritic spine remodeling rate increases at injury sites within the first two weeks following injury before decreasing (100), suggesting a way for damaged networks to increase variability and restore function. Furthermore, neuromodulation can stabilize disrupted network activity to promote network recovery (99). In the crab, photoinactiving LPG or blocking LG, IC, and DG glutamatergic synapses via picrotoxin could be viewed as injury states, where the functional synapses left behind are sufficient to maintain a coordinated pattern among gastric mill CPG neurons. Thus, in addition to multiple solutions to carry out behavior, synaptic degeneracy may also enable networks to compensate for loss of function following injury.

## Conclusion

Here, we found that although LPG is not necessary for gastric mill rhythm generation in the Gly^1^-SIFamide modulatory state, it contributes to pattern generation. Specifically, LPG regulates the activity level of other gastric mill neurons in our typical in vitro conditions. Additionally, we found that in the absence of glutamatergic chemical transmission, LPG slow bursts are sufficient to coordinate the remainder of the gastric mill neurons. This suggests degeneracy in synaptic connectivity and the possibility that there is a distinct behavioral or modulatory state in which LPG plays a dominant role in coordinating gastric mill neurons. Overall, we found that a switching neuron that is recruited to oscillate at a second network frequency can play an active role in the second network. This contrasts with a passive role for switching neurons recruited via internetwork synapses. It will be interesting to examine other instances of intrinsic property modulation as a mechanism of recruiting a neuron into another network to determine if this dichotomy between modulation of intrinsic and synaptic properties is a ubiquitous theme among neuronal networks.

## Acknowledgements

We thank Logan Fickling for help with spike phase analysis. This work funded by National Science Foundation (IOS: 1755283; DMB) and the Biology Department at Miami University.

---

**Supplemental Figure S1.**
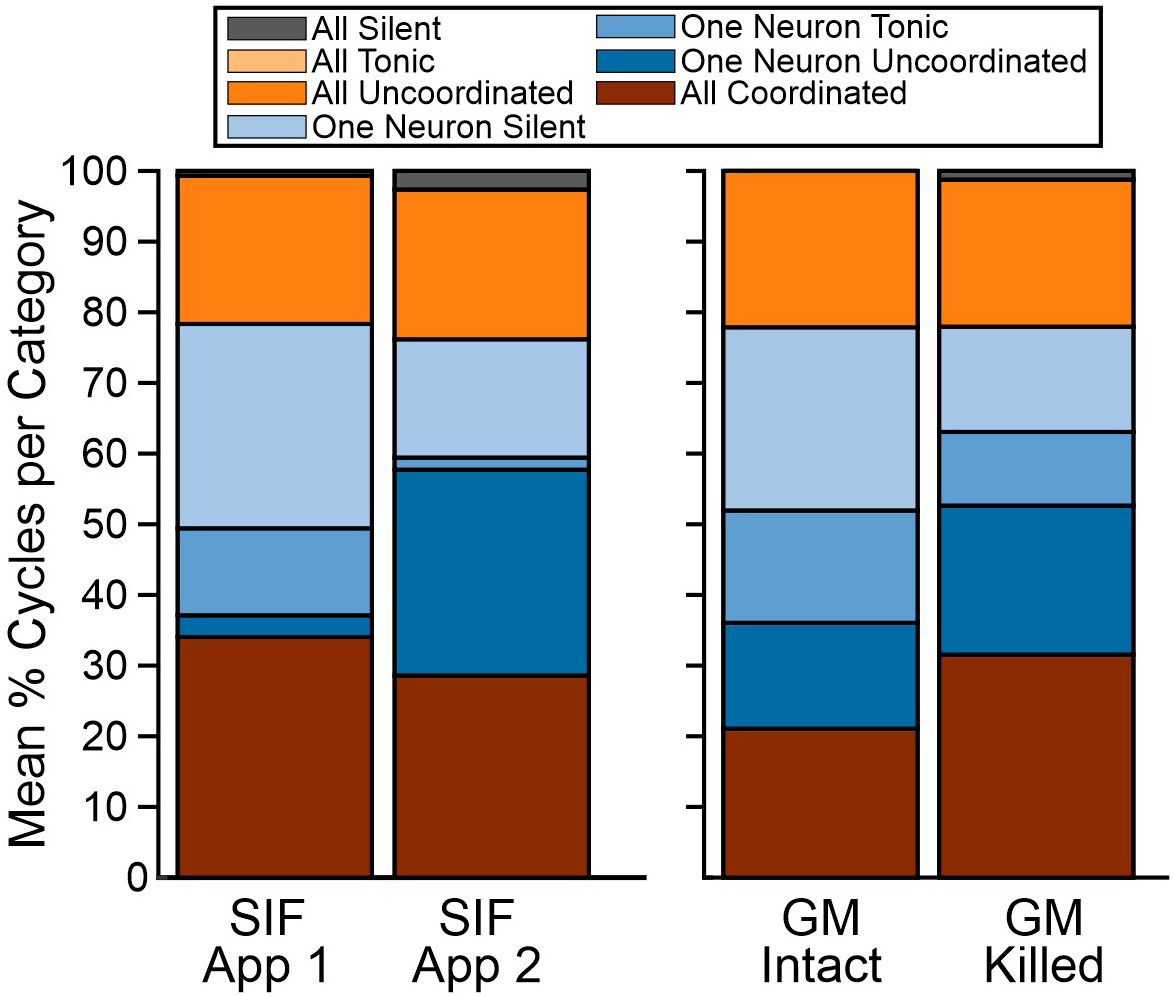
Stacked bar graphs of gastric mill network categories for LG, IC, LPG, and DG neurons during two consecutive Gly^1^-SIFamide applications (SIF App 1 versus SIF App 2, *left*), and during Gly^1^-SIFamide applications with GM Intact versus GM Killed (*right*).

**Supplemental Table S1.** Mean ± SEM percentage of identified gastric mill categories during two consecutive Gly^1^-SIFamide applications (SIF App)

| Coordination<br>Category | SIF App 1 | SIF App 2 | *t/z* | *p* |
| --- | --- | --- | --- | --- |
| All Coord. | 34.05 ± 9.49 | 28.61 ± 3.40 | 0.653* | 0.55 |
| All Uncoord. | 20.98 ± 7.12 | 21.20 ± 5.52 | -0.051* | 0.962 |
| One Uncoord. | 3.07 ± 2.07 | 29.12 ± 12.24 | -2.378* | 0.077 |
| One Tonic | 12.32 ± 5.16 | 1.71 ± 1.31 | 1.771* | 0.151 |
| One Silent | 28.91 ± 5.17 | 16.75 ± 6.04 | 1.572* | 0.191 |
| All Tonic | 0 | 0 | 1.267E-250** | 1 |
| All Silent | 0.68 ± 0.49 | 2.61 ± 1.50 | -1.784* | 0.149 |

Coordination categories identified in Gly<sup>1</sup>-SIFamide, \*t-statistic (t), paired t-test, \*\*z-statistic (z), Wilcoxon Signed Rank test

**Supplemental Table S2.** Mean ± SEM percentage of identified gastric mill categories during Gly^1^-SIFamide application with GM Intact versus GM Killed.

| Coordination |  |  |  |  |
| --- | --- | --- | --- | --- |
| Category | GM Intact | GM Killed | *t/z* | *p* |
| All Coord. | 21.08 ± 8.43 | 31.56 ± 6.56 | -1.053* | 0.333 |
| All Uncoord. | 22.11 ± 2.98 | 20.82 ± 3.92 | 0.297* | 0.777 |
| One Uncoord. | 14.97 ± 9.41 | 21.09 ± 8.31 | -0.65* | 0.54 |
| One Tonic | 15.91 ± 10.08 | 10.42 ± 6.71 | 0.6* | 0.569 |
| One Silent | 25.93 ± 7.81 | 14.90 ± 3.98 | 1.418* | 0.206 |
| All Tonic | 0 | 0 | 1.552E-251** | 1.000 |
| All Silent | 0 | 1.21 ± 0.80 | 1.342** | 0.5 |

Coordination categories identified in Gly<sup>1</sup>-SIFamide, \*t-statistic (t), paired t-test, \*\*z-statistic (z), Wilcoxon Signed Rank test

## Notes

### Competing Interest Statement

The authors have declared no competing interest.

